# Cyclophilin A regulates protein phase separation and mitigates haematopoietic stem cell aging

**DOI:** 10.1101/2021.02.24.432737

**Authors:** Laure Maneix, Polina Iakova, Shannon E. Moree, Jordon C.K. King, David B. Sykes, Cedric T. Hill, Borja Saez, Eric Spooner, Daniela S. Krause, Ergun Sahin, Bradford C. Berk, David T. Scadden, André Catic

## Abstract

Loss of protein quality is a driving force of aging^1^. The accumulation of misfolded proteins represents a vulnerability for long-lived cells, such as haematopoietic stem cells. How these cells, which have the ability to reconstitute all haematopoietic lineages throughout life^2^, maintain their regenerative potential and avert the effects of aging is poorly understood. Here, we determined the protein content in haematopoietic stem and progenitor cells to identify prevalent chaperones that support proteome integrity. We identified Peptidyl-Prolyl Isomerase A (PPIA or Cyclophilin A) as the dominant cytosolic foldase in this cell population. Loss of PPIA accelerated aging in the mouse stem cell compartment. In an effort to define targets of PPIA, we found that RNA- and DNA-binding proteins are common substrates of this chaperone. These proteins are enriched in intrinsically disordered regions (IDRs), which can catalyse protein condensation^3^. Isomerized target prolines are almost exclusively located within IDRs. We discovered that over 20% of PPIA client proteins are known to participate in liquid-liquid phase separation, enabling the formation of supramolecular membrane-less organelles. Using the poly-A binding protein PABPC1 as an example, we demonstrate that PPIA promotes phase separation of ribonucleoprotein particles, thereby increasing cellular stress resistance. Haematopoietic stem cell aging is associated with a decreased expression of PPIA and reduced synthesis of intrinsically disordered proteins. Our findings link the ubiquitously expressed chaperone PPIA to phase transition and identify macromolecular condensation as a potential determinant of the aging process in haematopoietic stem cells.

## Main Research Findings

Haematopoiesis is a dynamic regenerative process. Haematopoietic stem cells (HSCs) give rise to rapidly dividing progenitor cells that spawn hundreds of billions of progeny cells on a daily basis^2^. In contrast to progenitor cells, stem cells are long-lived, highly durable, and have a low mitotic index. Given their longevity and the absence of frequent cell division as a means of disposing of protein aggregates, protein homeostasis (proteostasis) is a critical component of HSC biology. Several studies have shown that proteostasis is tightly controlled in HSCs^4–6^ and the capacity of cells to maintain proteostasis declines with age^7^. Ensuring protein homeostasis requires precise control of translation, protein folding, transport, and degradation. Supporting enzymes (chaperones) are key actors in this network, ensuring efficient folding of newly translated polypeptides and conformational maintenance of pre-existing proteins.

To identify the most prevalent chaperones in the haematopoietic compartment, we analysed the proteome of mouse haematopoietic stem and progenitor cells (HSPCs) through semi-quantitative 2-D gel electrophoresis and mass spectrometry (Fig. 1A and Extended Data Fig. 1). Of the discernible protein peaks, Cyclophilin A (PPIA) accounted for up to 14% of the cytosolic proteome, making it the most abundant foldase in HSPCs. PPIA was also the most highly expressed chaperone at the transcript level, accounting for over 0.5% of all mRNAs (Extended Data Fig. 2). Prolyl isomerases are conserved enzymes and cyclophilins represent one of the four classes with 17 members in humans^8^ (Extended Data Fig. 3). Cyclophilins catalyse the isomerization of proline, the only proteinogenic amino acid that exists in abundance in both *trans-* and *cis-*configurations^9^. Previously described PPIA knockout (PPIA^-/-^) mice^10^ demonstrated the gene is non-essential and animals showed no apparent phenotype under homeostatic conditions in the C57BL/6 background^11^.

**Figure 1:**
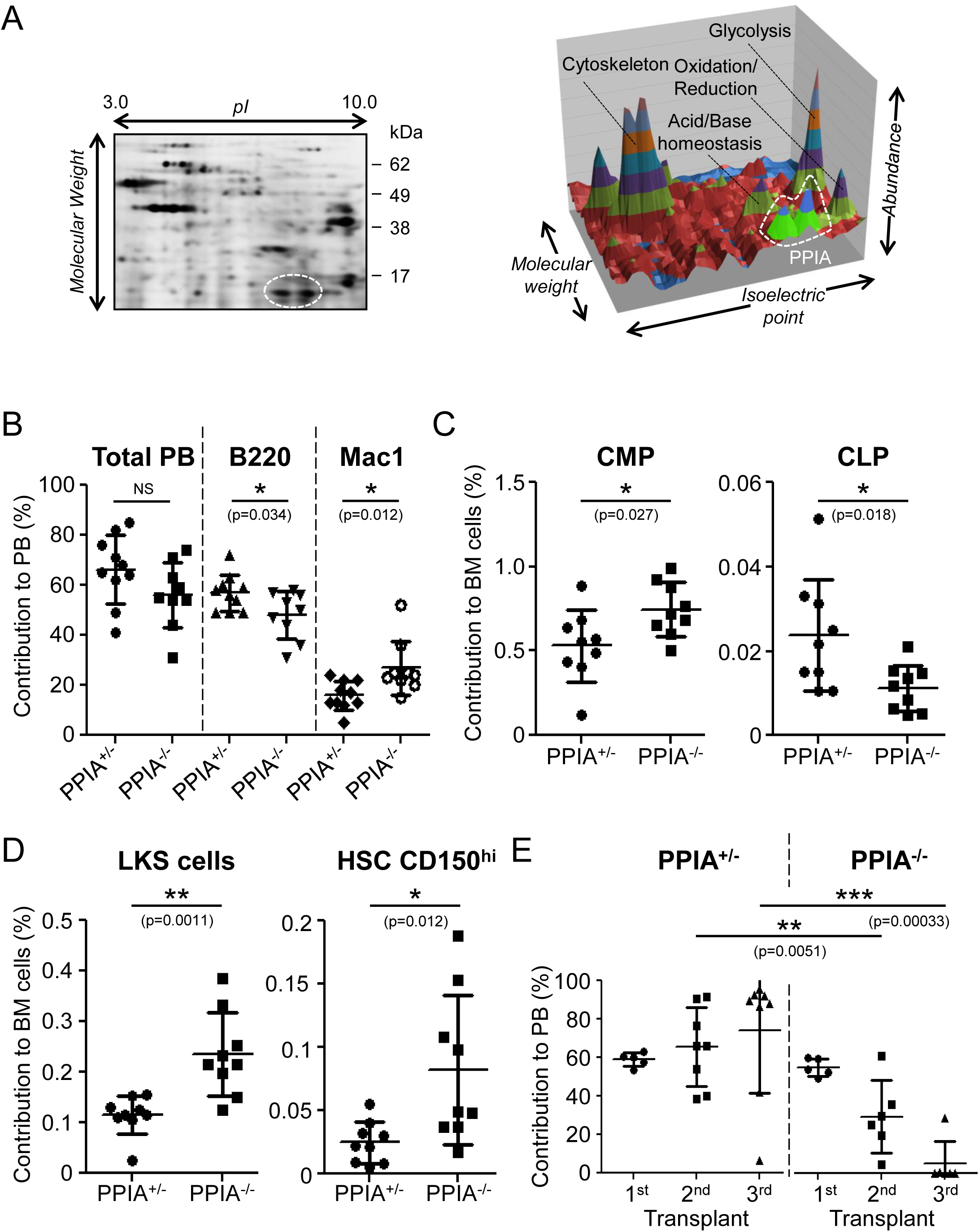
PPIA deficiency induces an aging-like haematopoietic phenotype. **A**, Left panel: Haematopoietic stem and progenitor cell (HSPC) lysate was labelled with amine- reactive dye and separated on a 2-D electrophoresis gel. Right panel: Quantitative intensity representation of the mouse HSPC proteome. Dashed white lines indicate PPIA as determined by MS/MS. Two independent experiments were performed. **B**, Six months after bone marrow (BM) transplantation, blood cell analysis with flow cytometry reveals that PPIA knockout (PPIA^-/-^) BM donor cells give rise to less lymphoid (B220) and more myeloid (Mac1) cells in the peripheral blood (PB). **C**, Seven months after transplantation, BM cell analysis with flow cytometry shows that mice transplanted with PPIA^-/-^ BM donor cells have increased common myeloid progenitors (CMP) and decreased common lymphoid progenitors (CLP). **D**, PPIA-deficient donor cells show increased HSPCs (LKS; lineage^-^/c-Kit^+^/Sca1^+^ cells) and CD150^high^ (lineage^-^/c-Kit^+^/Sca1^+^/CD34^-^/CD135^-^/CD150^high^) haematopoietic stem cells (HSCs). **E**, Cell analysis shows that PPIA^-/-^ donor-derived progenitor cells exhaust in serial transplantations. Shown is the proportion of donor-derived (CD45.2^+^) cells among all PB cells 2, 4, and 12 months after the first transplantation. For B, C, D, and E, data are mean ± SD; NS, non- significant, *p < 0.05, **p < 0.01, and ***p < 0.001 by unpaired Student’s two-tailed *t*-test. Data are representative of two independent biological replicates.

**Figure 2:**
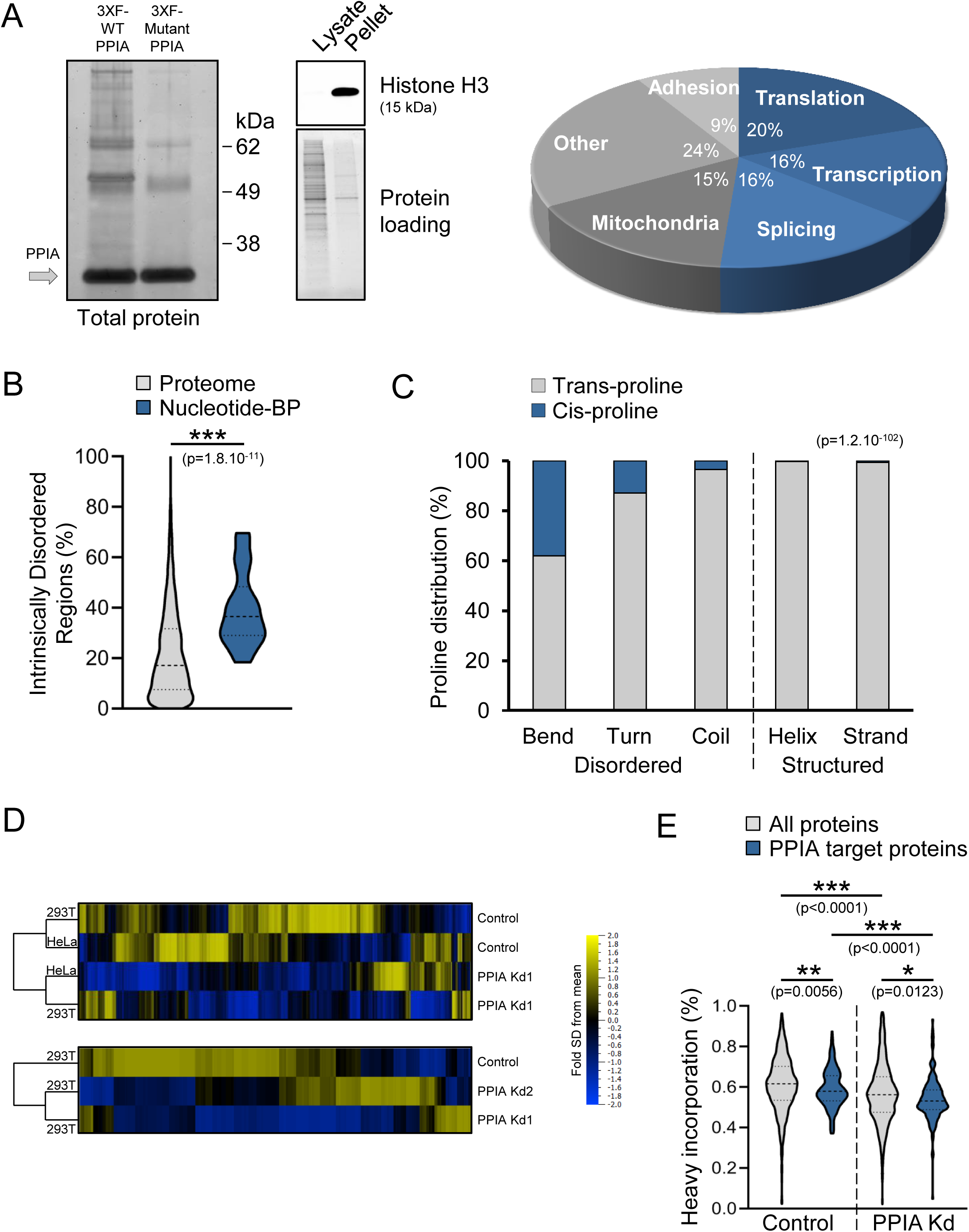
PPIA activity promotes translation of proteins enriched for Intrinsically Disordered Regions (IDRs). **A**, Left panel: Immunoprecipitation (IP) and SYPRO Ruby gel stain of triple FLAG-tagged wild- type (3XF-WT PPIA) and G104A point-mutant PPIA (3XF-Mutant PPIA) were performed in order to identify PPIA interacting proteins. Grey arrow indicates positive enrichment for PPIA protein in the pull-down fractions. Middle panel: Purity of the cell lysates was verified by the detection of histone H3 protein expression in subcellular fractions. 3-D pie chart: Gene ontology enrichment analysis of 3XF-WT PPIA versus 3XF-Mutant PPIA IP-mass spectrometry results. Data represent 385 consistently identified proteins in 293T cells from two independent biological replicates. **B**, Quantification of the percentage of intrinsically disordered regions in the total proteome versus 3XF-WT PPIA-interacting nucleotide-binding proteins (BP) with nucleotide-binding domains containing an alpha-beta plait structure (InterPro identifier IPR012677). ***p < 0.001 determined by Wilcoxon rank-sum test. **C**, Spatial distribution of proline residues within secondary protein structures. The distribution of *cis*-prolines in structured vs disordered regions was compared with the Chi Square test with Yates’ correction and is based on a meta-analysis of solved protein structures^25^. **D**, Pulsed SILAC was performed with protein extracts from control or PPIA knockdown (Kd) cell lines in order to measure newly synthetized proteins, as described in Extended Data Fig. 8. **E**, Uptake of heavy amino acids by control or PPIA Kd 293T cells was quantified following pulsed SILAC experiment. *p < 0.05, **p < 0.01, and ***p < 0.001 determined by Wilcoxon rank- sum test.

**Figure 3:**
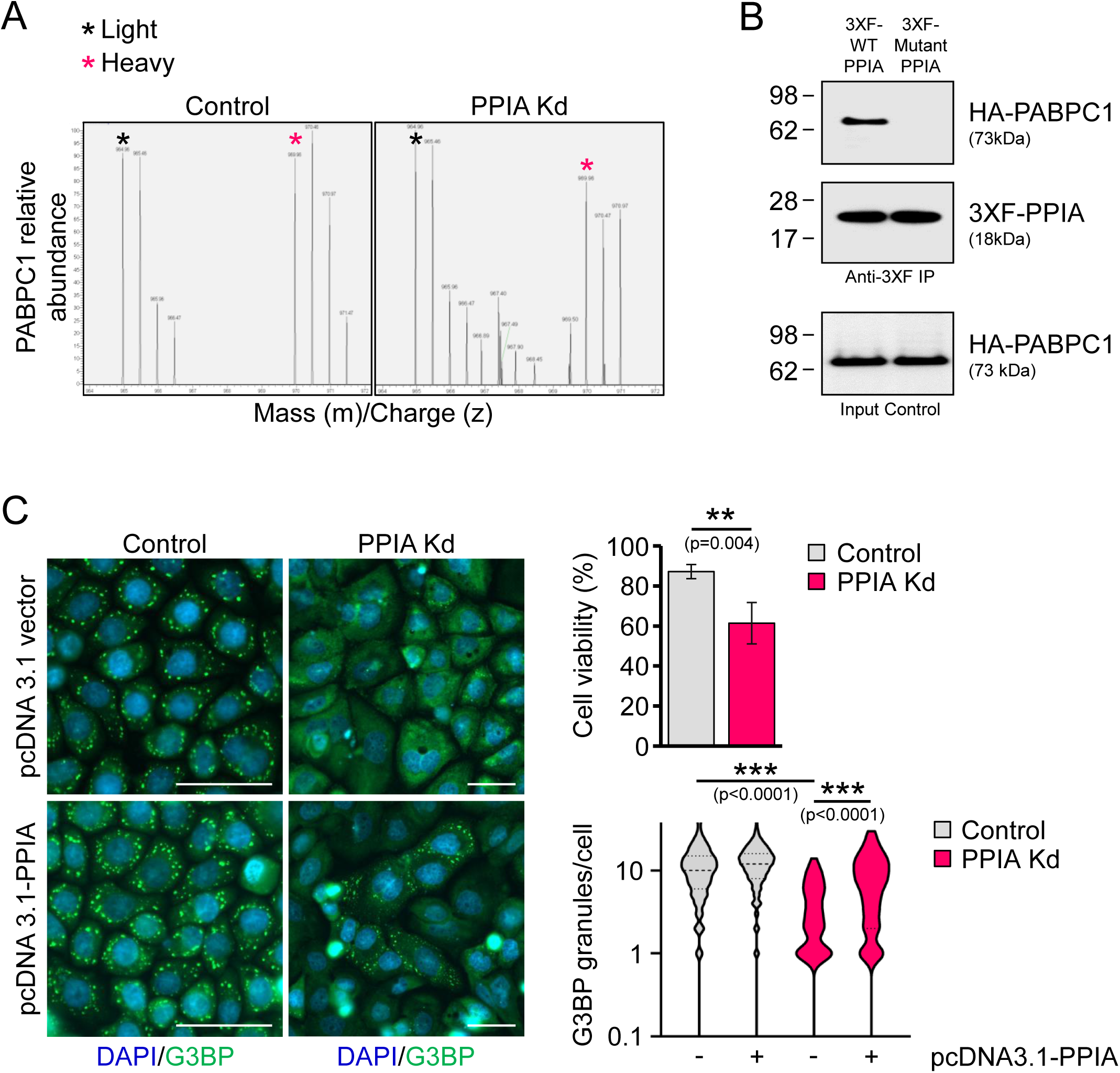
PPIA and its IDR-rich substrate PABPC1 regulate protein phase separation. **A**, Mass spectra for control and PPIA Kd HeLa cells show changes in PABPC1 protein turnover measured by pulsed SILAC. The asterisks denote the peaks for PABPC1 peptides. **B**, Immunoprecipitation of 3XF-WT PPIA and 3XF-Mutant PPIA in 293T cells was followed by Western blot analyses to detect poly(A)-binding protein 1 (PABPC1) and PPIA protein. The results were validated by more than three independent biological replicates. **C**, Stress granule formation was visualized and quantified with G3BP1 staining after stress induction with sodium arsenite in HeLa control or PPIA Kd cells. DAPI, blue; G3BP1, green; scale bar=50 µm. Fluorescence microscopy images are representative of three independent experiments. Data are mean ± SD of three replicate wells; **p < 0.01 by unpaired Student’s two- tailed *t-*test for cell viability measurements; ***p<0.001 by Wilcoxon rank-sum test for G3BP1 granule counting.

To assess the role of PPIA in haematopoiesis, we compared knockout and heterozygous animals in a series of functional assays after bone marrow transplants. In these assays, heterozygous animals were indistinguishable from wild types (data not shown). For these studies, we competitively transplanted CD45.2^+^ total nucleated bone marrow (BM) cells from PPIA^-/-^ or PPIA^+/-^ mice together with equal numbers of CD45.1^+^ wild-type support BM cells into lethally irradiated CD45.1^+^ recipient animals. Six months after transplantation, when the BM was fully repopulated by long-lived donor HSCs, we observed a statistically significant decrease of PPIA^-/-^ B lymphocytes in the blood of recipient animals. In contrast, myeloid cells were increased in recipients of knockout cells (Fig. 1B). Changes in BM progenitor cells drove this myeloid skewing in the peripheral blood, as we observed an increase in common myeloid PPIA^-/-^ progenitor cells at the expense of lymphoid progenitor cells (Fig. 1C). We also found higher relative and absolute numbers of HSPCs and myeloid-biased CD150^high^ HSCs^12^ in recipients of PPIA^-/-^ BM (Fig. 1D).

To functionally define stem cell activity, we performed limiting dilution transplantation experiments with PPIA^-/-^ and PPIA^+/-^ BM cells. The results were comparable, indicating that higher numbers of immunophenotypic stem cells in the knockout BM did not correlate with increased stem cell activity (Extended Data Fig. 4). Next, we tested the self-renewing ability of HSCs by measuring the repopulation of BM following serial transplantations of PPIA^-/-^ or PPIA^+/-^ donor cells. Unlike wild-type or heterozygous BM cells, PPIA^-/-^ cells failed to functionally engraft after the first round of transplantation and displayed exhaustion in long-term repopulation assays (Fig. 1E). Taken together, these functional transplant assays revealed cell- intrinsic defects leading to myeloid skewing, an immunophenotypic but not functional increase in stem cells, and impaired self-renewal with accelerated exhaustion in PPIA^-/-^ HSCs. These three characteristics are hallmarks of haematopoietic aging^13–16^, suggesting that the absence of PPIA resembles premature aging at the stem cell level.

**Figure 4:**
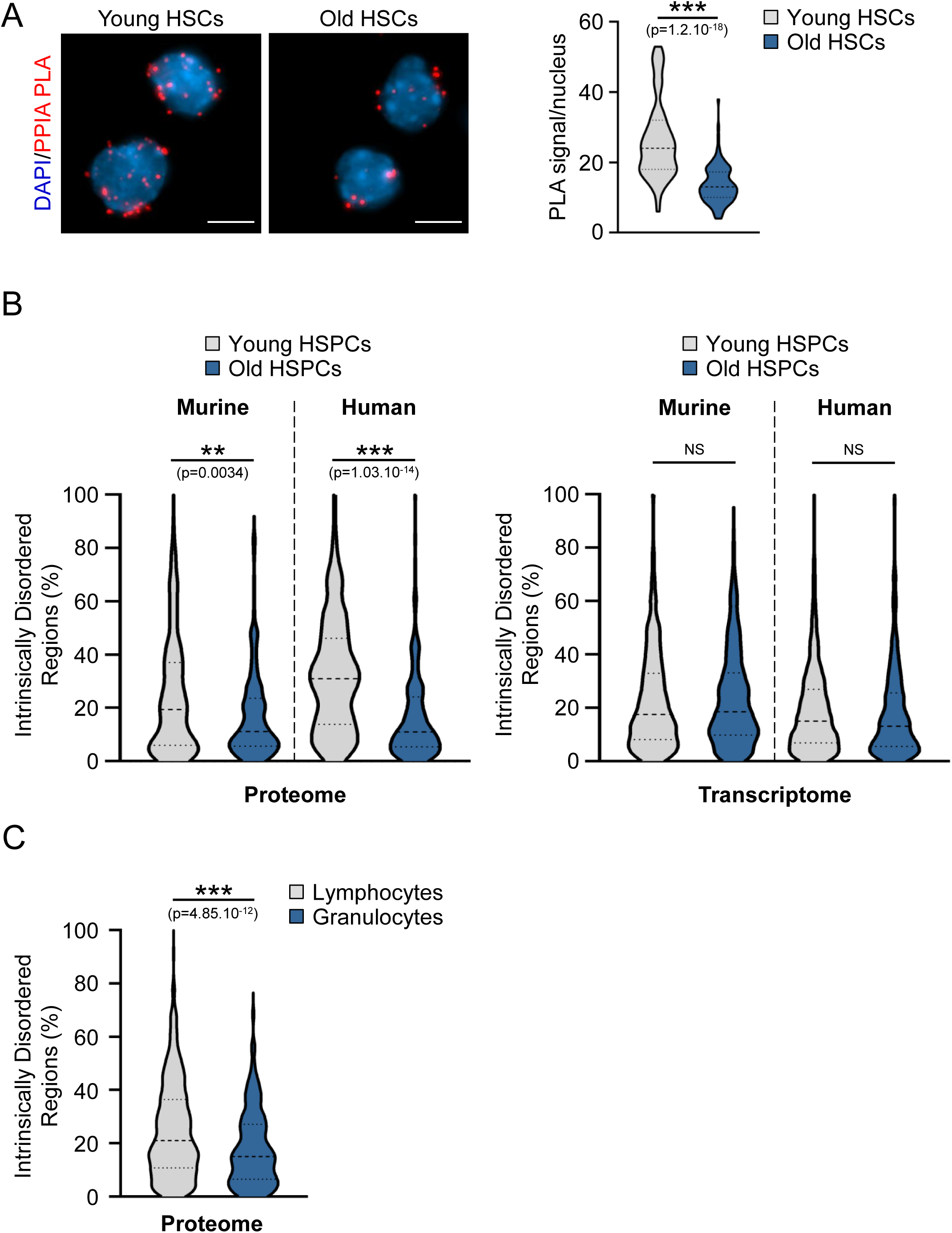
PPIA levels decline with age and contribute to IDR protein deficiency in HSPCs. **A**, Proximity Ligation Assay (PLA) to quantify PPIA protein levels in mouse HSCs shows decreased PPIA expression in 23 month-old HSCs when compared to 5 month-old cells. Fluorescence microscopy images are representative of two independent biological replicates. DAPI, blue; PPIA, red; scale bar=5 µm. ***p<0.001 by unpaired Student’s two-tailed *t*-test for quantification of PLA positive spots. **B**, Quantification of intrinsically disordered region content in the mouse and human proteome (left panel) and transcriptome (right panel). Analysis of murine RNA-seq and MS/MS data was validated by a total of two independent experiments. **p < 0.01 and ***p < 0.001 determined by Wilcoxon rank-sum test; NS, non-significant. **C**, Quantification of the percentage of intrinsically disordered regions in the unique protein content of human lymphocytes versus granulocytes. ***p < 0.001 determined by Wilcoxon rank- sum test.

PPIA is a highly and ubiquitously expressed prolyl isomerase that interacts with a wide range of proteins^17, 18^. PPIA isomerizes proline residues of *de novo* synthesized proteins during translation. Several *in vitro* refolding studies demonstrated that cis/trans-isomerization of prolyl bonds can be a rate-limiting step during normal protein folding and translation^19–22^. To gain insight into the molecular changes caused by its depletion, we sought to determine the client proteins of PPIA. We accounted for non-specific interactions and distinguished PPIA-specific substrate proteins using a previously identified PPIA G104A mutant^23^, which has reduced activity. We tested several mutations and found that inactivation of the catalytic core yielded insoluble PPIA, while the G104A mutation, which moderately reduces substrate access to the catalytic core trough an obstruent methyl group, allowed for normal expression levels and intracellular distribution (Extended Data Fig. 5). Therefore, proteins that interact with the wild-type PPIA but fail to bind to the PPIA G104A mutant are likely direct substrates of the foldase. Differential co-immunoprecipitation between the wild-type PPIA and the mutant PPIA revealed approximately 400 substrates of the wild-type enzyme (Fig. 2A, Extended Data Fig. 6, and SI-PPIA client proteins), including 39 transcriptional regulators (Extended Data Table 1). Since we performed the co-immunoprecipitation in the cytosolic cell fraction, these results suggest PPIA and substrates interact during translation and prior to nuclear translocation (Fig. 2A).

**Figure 5:**
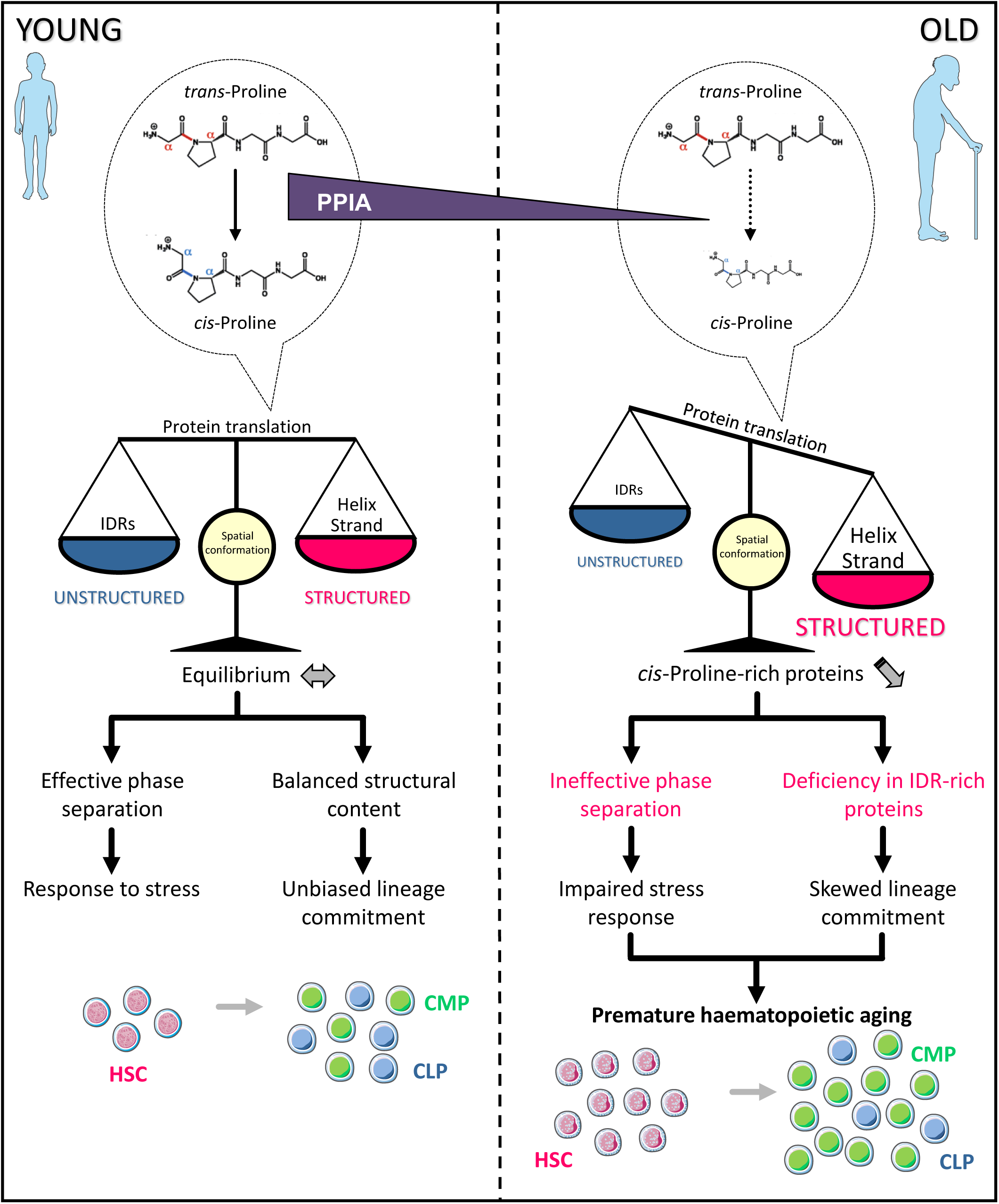
Model of PPIA activity and function in the aging haematopoietic compartment. PPIA supports nascent proteins during translation and affects *cis*-proline isomerization in IDRs. Therefore, proteins rich in IDRs require more foldase activity. IDRs are essential for phase separation and also more highly expressed in the lymphoid compared to myeloid lineages. PPIA expression decreases during haematopoietic aging and the aged proteome is consequently depleted of proteins involved in phase separation and hampered in the production and function of lymphocytes. In conclusion, PPIA deficiency impairs stress response in HSCs, biases lineage commitment, and accelerates HSC aging.

In addition to DNA-binding proteins, the most prevalent group of PPIA clients included RNA-associated proteins involved in ribosome assembly, translation, and splicing (Fig. 2A). Among structural features that were enriched in PPIA substrates, the most significant motif was the nucleotide-binding domain with an alpha-beta plait structure. Proteins containing these domains also feature high levels of intrinsically disordered regions (IDRs), *i.e.* unstructured protein regions displaying a sequence-driven preference for conformational heterogeneity^24^ (Fig. 2B).

Prolyl isomerases such as PPIA catalyse the reversible *trans-* and *cis-*conversions of peptide bonds at proline residues. While *cis-*prolines represent about 10% of the total proline pool, they are disproportionally enriched within IDRs (Fig. 2C), based on an analysis of over 15,000 proline residues within the Protein Data Bank^25^. Thus, these findings indicate that PPIA and related enzymes predominantly isomerize prolines within unstructured protein regions.

In line with PPIA’s proposed activity as a co-translational chaperone^26^ and given that proline isomerization is slow and often rate-limiting during translation, we expected PPIA expression to directly affect *de novo* protein translation of its substrates. We determined the *de novo* synthesis of proteins using pulsed SILAC in HeLa and 293T cells with either normal or reduced levels of PPIA (Extended Data Fig. 7 & Extended Data Fig. 8). We found that loss of PPIA significantly reduced translation, specifically of PPIA clients, in both cell types (Fig. 2D and SI-Pulsed SILAC). Overall, *de novo* translation rates of PPIA target proteins were lower than for other proteins and further reduced when PPIA was depleted (Fig. 2E). These results demonstrate that PPIA supports *de novo* translation of its target proteins and are consistent with earlier reports that IDR-rich proteins have a low translation rate, *i.e.* synthesis of disordered proteins appears to be at a disadvantage^27, 28^ (Extended Data Fig. 9).

Within PPIA substrate proteins, the dominant ontologies we found were translation and mRNA splicing, and we discovered that more than 20% of PPIA clients are known to engage in protein phase separation (reviewed in^29–33^) (Extended Data Table 2). Liquid-liquid phase separation allows the intracellular compartmentalization of ribonucleoprotein assemblies through changes in solubility and subsequent formation of membrane-less organelles^34^. Prominent examples of phase-separating proteins that bind to wild-type but not mutant PPIA include the splicing factor TDP43 (TARDBP), the nucleolus organizing transcription factor UBTF, the stress granule initiator G3BP1, DEAD-box helicases, YBX1, and the mRNA regulator Poly(A) Binding Protein Cytoplasmic 1 (PABPC1). Recently, the notion has emerged that intrinsically disordered polypeptides can be molecular determinants of phase separation and modulate the formation of membrane-less organelles^35–37^. For instance, the RNA binding protein PABPC1 has been shown to engage in phase separation through its unstructured proline-rich linker region, which is instrumental to the formation of RNA stress granules^38^. Following PPIA knockdown, *de novo* synthesis of PABPC1 protein was reduced by 20-30%, suggesting the foldase engages with PABPC1 during translation (Fig. 3A). We biochemically confirmed that PABPC1 is a client of PPIA (Fig. 3B; Extended Data Fig. 6 and Extended Data Fig. 10) and that PABPC1 protein expression is reduced following PPIA inhibition by cyclosporine A (Extended Data Fig. 11). In contrast, PPIA inhibition did not reduce the transcription of the gene encoding PABPC1 (Extended Data Fig. 11). These findings support the notion that PPIA regulates PABPC1 folding by controlling proline isomerization during translation.

In response to diverse stresses or unfavourable growth conditions, PABPC1 undergoes phase transition and participates in stress granule formation to sequester cytoplasmic RNA and ribosomes (SI-Video 1). This allows cells to temporarily reduce protein translation^39, 40^. Treating cells with the oxidative stressor sodium arsenite is a well-established experimental approach to study formation of stress granules. To determine whether proline isomerization affects phase separation of PABPC1, we genetically modulated PPIA activity and assessed the dynamics of stress granule formation. Upon stress induction with sodium arsenite, we observed significantly reduced stress granule formation in the absence of prolyl isomerase activity (Fig. 3C) (SI-Videos 2 and 3). Cells devoid of PPIA were more susceptible to cell death following oxidative stress (Fig. 3C). In addition, re-introduction of the foldase partially rescued stress granule formation in PPIA knockdown cells (Fig. 3C).

Our data suggest that PPIA is involved in PABPC1 folding during translation and that the foldase plays a critical role in phase transition of this protein. Given the number of key regulators of phase separation among PPIA clients, our findings indicate that proline isomerization may be a major determinant for the formation of membrane-less organelles. Supporting this hypothesis, PPIA has been shown to co-localize with stress granules^41^. In addition, proline residues are key residues during phase transition of prion-like and unstructured proteins, and can regulate protein solubility and amyloid formation in an isomer-specific fashion^34, 42–45^.

We next addressed whether our findings that PPIA regulates the function of intrinsically disordered polypeptides, and thereby protein phase separation, relate to the haematopoietic phenotype resembling aging that we observed in PPIA^-/-^ mice. A previous quantitative proteomic analysis of human dermal fibroblasts from subjects of different ages showed that PPIA is significantly reduced with age, but its role in haematopoietic aging remained unknown^46^. In the haematopoietic compartment, we found a substantial reduction of PPIA protein in HSCs from old mice compared to younger cells (Fig. 4A). Of note, PPIA transcripts were not altered in HSPCs of different ages in our RNA-seq data. Based on our earlier findings, we would expect reduced PPIA activity to result in lower expression of IDR-rich proteins. Indeed, the comparative analysis of the proteome of young and old mice showed a reduction of IDR-rich proteins in HSPCs during aging, which is not reflected at the transcriptome level (Fig. 4B). These results suggest that synthesis of IDR-rich proteins, which are challenging to translate, is reduced in older HSCs and progenitor cells, and may therefore affect protein phase separation and stress responses in these cells.

We next explored whether our findings in the mouse haematopoietic system also applied to human blood. A comprehensive study of the HSPC transcriptome and proteome in humans of different ages^47^ confirms our results and shows highly significant reduction in the expression of intrinsically disordered proteins in aged individuals that is not manifest in the transcriptome (Fig. 4B). Given the disconnect we observed between protein and gene expression, we speculate that translation efficiency may be partially responsible for the differences in lineage commitment between young and old haematopoiesis, namely the shift towards myelopoiesis at the expense of lymphopoiesis during aging. Intrinsically disordered proteins are less efficiently translated, in part due to their dependence on foldases such as PPIA. At the same time, these proteins are under positive selection during the course of evolution and complex organisms contain more disordered regions in their proteomes^48, 49^. We hypothesized that this phylogenetic feature makes IDR-rich proteins more prevalent in evolutionarily newer lymphoid cells compared to older myeloid cells. Indeed, the proteome of human granulocytes showed significantly fewer disordered regions compared to lymphocytes (Fig. 4C). As a consequence, the proteome of granulocytes is likely more efficiently synthesized, which may partially explain the preferred generation of these cells over lymphocytes with age or under stress. We therefore propose that the expression of IDR-rich proteins constitutes a translational bottleneck during aging or under stress conditions.

In conclusion, we discovered that Cyclophilin A (PPIA) is the most highly expressed chaperone in haematopoietic stem and progenitor cells. PPIA engages with its substrate proteins during translation and our data suggests that it does so largely through *cis-*isomerization of prolines within intrinsically disordered regions. A substantial fraction of PPIA targets engage in phase separation and we found evidence for impaired protein condensation in the absence of the foldase. PPIA levels are reduced in aged haematopoietic stem cells, altering the proteome and thereby decreasing stress resistance, reducing the long-term fitness of these cells, and biasing their lineage commitment. Altogether, our results identify PPIA as a major component in the chain of molecular events during haematopoietic aging that affects the expression of disordered proteins and the formation of membrane-less organelles (Fig. 5).

## Supporting information

SI-Mouse HSPCs RNA seq

SI-PPIA client proteins

SI-Pulsed SILAC

## Methods

### 2-D electrophoresis gels

We used the Zoom IPGRunner system (Invitrogen) to separate proteins in two dimensions. We isolated HSPCs of young animals (4-8 months of age) and lysed them according to the manufacturer’s instructions with urea, CHAPS, DTT, and ampholytes. CyDyes DIGE fluors (minimal dyes) were used according to the vendor (Amersham) with fluorophores Cy3 and Cy5 for post-lysis labelling to ensure that only 1-2% of lysines are labelled in a given protein. Labelling intensities were measured with a Typhoon FLA 9000 scanner and quantified with DeCyder 7.0 and ImageQuant software (GE Healthcare). We normalized total protein abundance based on protein size and lysine concentration for spots with known identity by MS/MS. PPIA quantity was identical in two independent experiments using different dyes.

### Tandem mass spectrometry (MS/MS)

#### Characterization of the mouse HSPC proteome following 2-D electrophoresis

Following trypsinolysis, we analysed digested peptides by reverse-phase liquid chromatography electrospray ionization mass spectrometry using a Waters NANO-ACQUITY-UPLC, coupled to a Thermo LTQ linear ion-trap mass spectrometer. In order to identify proteins, we searched MS/MS spectra against the non-redundant NCBI protein database using SEQUEST program (http://proteomicswiki.com/wiki/index.php/SEQUEST).

#### PPIA protein complex identification following 3XF-PPIA immunoprecipitation

Immunopurified samples were analysed by mass spectrometer-based proteomics as previously described^50^. Minor modifications from the previously cited protocol are listed here. Digested peptides were injected into a nano-HPLC 1000 system (Thermo Scientific) coupled to LTQ Orbitrap Elite (Thermo Scientific) for first repeat and Q Exactive Plus (Thermo Scientific) mass spectrometer for second repeat samples. Separated peptides were directly electro-sprayed into the mass spectrometer, controlled by the Xcalibur software (Thermo Scientific) in data-dependent acquisition mode, selecting fragmentation spectra of the top 25 and 35 strongest ions for 1^st^ samples and 2^nd^ samples, respectively. MS/MS spectra were searched against target-decoy human RefSeq database (released January 2019, containing 73637 entries) with the software interface and search parameters previously described^50^. Variable modification of methionine oxidation and lysine acetylation was allowed. Protein abundance was calculated with the iBAQ algorithm and relative protein amounts between samples were compared with an in-house processing algorithm^51^.

#### Mouse HSPC global proteome profiling

Whole bone marrow (BM) was isolated from the femurs and tibias of 3-month old and 21 month-old wild-type male mice. Magnetic depletion of lineage-positive haematopoietic cells was performed using the EasySep mouse haematopoietic progenitor cell enrichment kit (Stem Cell Technologies), and lineage-depleted stem and progenitor cells were submitted to mass spectrometry analysis. Following sample lysis and overnight trypsin digestion, reconstituted peptidic fractions were loaded onto a nano-HPLC 1000 system (Thermo Fisher Scientific) coupled to an Orbitrap Fusion Lumos Tribrid mass spectrometer (Thermo Fisher Scientific), with identical acquisition settings as previously described^52^. Trap and capillary HPLC columns have been previously described^50^. The search of resultant MS/MS spectra against target-decoy mouse RefSeq database (released June 2015, containing 58549 entries) was done in Proteome Discoverer 2.1 interface (Thermo Fisher) with Mascot 2.4 algorithm (Matrix Science). Variable modifications allowed were methionine oxidation and protein N-terminal acetylation. Search settings were the following: precursor mass tolerance of 20 ppm, a maximum of two missed trypsin cleavages, and fragment ion mass tolerance of 0.5 Dalton. Assigned peptides were filtered with 1% false discovery rate FDR. The in-house iFOT data processing algorithm^51^ was used to calculated label free relative abundance of proteins in samples.

GO analyses were performed with the DAVID bioinformatic database (https://david.ncifcrf.gov). The degree of native protein disorder was determined using the openly available large-scale DisoDB database (http://bioinfadmin.cs.ucl.ac.uk/disodb/) ^28^ DisoDB is based on an algorithm that predicts intrinsically unstructured segments and secondary structures in specific proteins with a confidence score, using available data from the Ensembl website (v.48). A disordered region was defined as a protein segment having at least 30 contiguous disordered residues. Additionally, spatial conformation of proline residues was calculated based on previously published structures and with respect to the peptide bond conformation (*cis* or *trans*)^25^.

### Transplantations

C57BL/6 mice were lethally irradiated with a Cs137 source at a single dose of 9.5 Gy up to 24 hours prior to transplantation.

#### Peripheral blood (PB) and bone marrow (BM) cell analysis

Cells were injected into the tail vein in 100 µl PBS. 375,000 nucleated bone marrow cells of PPIA heterozygous (PPIA^+/-^) or knockout (PPIA^-/-^) mice (CD45.2^+^) were co-injected with the same number of CD45.1^+^ competitor cells. We analysed PB at weeks 5, 8, 12, and 24 (shown are the 24 week analyses). Trend wise differences between PPIA knockout and heterozygous cells emerged at weeks 8 and 12 (p<0.1) and became statistically significant at the 24 week analysis. Final BM collection occurred at week 28.

#### Serial transplantations

Equal numbers of BM cells (500,000 cells) from donor mice were mixed with 500,000 competitor cells from wild-type mice and injected into lethally irradiated recipient mice. Two months after primary transplantation, 1,000,000 nucleated BM cells from the primary recipients were harvested for a second and again after 2 months for a third round of transplantation. Final evaluation was performed 6 months after the third transplantation. All donor and recipient animals were gender-matched and between 3-6 months of age. Separate experiments were conducted in male and female mice with identical results. Experiments had a statistical power of >90% and final results were based on at least five animals per group. Transplant recipient animals were randomly assigned at the time of irradiation and donor cells were pooled from up to three animals. PPIA^+/-^ animals and cells were indistinguishable from wild-type animals and cells in all experiments tested (data not shown). PPIA^+/-^ mice were generated by backcrossing into C57BL/6J for over ten generations.

### Mice

PPIA^+/-^, PPIA^-/-,^ and C57BL/6 wild-type animals used in transplantation studies were kept at the Massachusetts General Hospital (MGH) Simches facility under pathogen-free conditions and treated according to MGH Institutional Animal Care and Use Committee (IACUC)-approved protocols. PPIA^-/-^ mice were born at sub-Mendelian ratios but displayed no abnormal phenotype after multiple generations of backcrossing to the C57BL/6 genetic background. All procedures on mice were performed with approval from the MGH and Baylor College of Medicine IACUC and followed guidelines from the National Institutes of Health Guide for the Care and Use of Laboratory Animals. All mice were housed in ventilated cages, on a standard rodent diet of chow and water *ad libitum*, under a 12-hour light/dark cycle. Animals with signs of sickness or infection were excluded from the study.

### Cell analysis and FACS

First, freshly isolated PB and BM were initially depleted of lineage positive cells with MACS LD columns (Miltenyi Biotec), as previously described^53^. Then, cells were analysed with an LSR II instrument and isolated with an Aria I fluorescence-activated cell sorter (BD Biosciences).

The following antibody combinations were used for cell phenotyping: HSPC (c-Kit^+^, lineage^-^), LKS (c-Kit^+^, Sca1^+^, lineage^-^), CMP (c-Kit^+^, Sca1^-^, lineage^-^, CD16/32^-^, CD34^+^), CLP (c-Kit^int.^, Sca1^int.^, lineage^-^, CD127^+^, CD34^+^), HSC (c-Kit^+^, Sca1^+^, lineage^-^, CD135^-^, CD34^-^, CD150^+^). Immunostainings were performed by incubating cells with anti-c-Kit (clone 2B8, BD Biosciences or Life Technologies), anti-Sca1 (clone D7, Caltag Medystems or Thermo Fisher Scientific), anti-CD16/32 (clone 93, eBioscience), anti-CD34 (clone RAM34, BD Biosciences), anti-CD135 (clone A2F10.1, BD Biosciences), anti-CD150 (clone TC15-12F12.2, BioLegend), anti-CD127 (clone SB/199, BioLegend) and anti-CD45.1/2 (clones A20 and 104, BioLegend) antibodies for 30 min (PB) or 60 min (BM) at 4°C prior to FACS analyses.

The antibodies used for lineage depletion were anti-CD11b (clone M1/70, BD Biosciences), anti-Ly-6G and Ly-6C (clone RB6-8C5, BD Biosciences), anti-CD8α (clone 53- 6.7, BD Biosciences), anti-CD3ε (clone 145-2C11, BD Biosciences), anti-CD4 (clone GK1.5, BD Biosciences), anti-TER-199 (clone TER-119, BD Biosciences), anti-CD45R (clone RA3- 6B2, BD Biosciences) and streptavidin (S32365, Thermo Fisher Scientific). The source of the samples was blinded to the FACS analyst.

### Cell culture and Drug treatments

Biochemical assays were performed in 293T or HeLa cells which were maintained at 37 °C in a humidified incubator containing 5% CO_2_. Cell lines were purchased from ATCC, cultured with the medium composition recommended by the supplier, and monitored for signs of infection, including mycoplasma contamination.

Stable 293T or HeLa control and PPIA Kd1/Kd2 cell lines were generated using pLKO.1 lentiviral vectors encoding short hairpin RNAs targeting the human PPIA protein (clone ID# TRCN0000049171 (Kd1) or clone ID# TRCN0000049170 (Kd2), Horizon Discovery) designed by The RNAi Consortium (TRC). Cell lines stably transduced with a pLKO.1 TRC empty vector encoding a non-targeting sequence (clone ID# TRC TRCN0000241922, Horizon Discovery) served as controls. Following puromycin selection (2 µg/ml, Gibco, Fisher Scientific), PPIA knockdown efficiency was assessed by measuring PPIA protein expression by Western blots in stably transduced cells (Extended Data Fig. 7). The two constructs PPIA Kd1 and PPIA Kd2 showed >80% knockdown efficiency by immunoblot and were tested independently.

Control and PPIA Kd1 Hela cells were transfected with pcDNA3.1-PPIA vector or corresponding pcDNA3.1 control vector for 48h. Following stress induction with sodium arsenite (50 µM, Sigma Aldrich) for 1h, immunostaining for G3BP1 protein, a marker of stress granule assembly, was performed using a rabbit polyclonal anti-G3BP1 antibody (Cat. #13057- 2-AP, Proteintech). The cells were mounted by Prolong gold antifade mounting medium containing DAPI (Invitrogen) and were imaged at 20X magnification on a Celldiscoverer 7 confocal microscope (Zeiss) operated with the Zen pro imaging software (Zeiss). The exposure time and gain were maintained at a constant level for all samples, and the stress granule analysis was carried out with the ImageJ software. Cell viability was measured on a Cellometer Auto 2000 automated cell counter with the ViaStain AO/PI staining solution (Nexcelom Bioscience).

Fig. 1, 4A, and 4B are based on freshly isolated murine HSPCs and HSCs. Fig. 2A, 2B, 2C, 2D, 2E, and 3B show experiments with 293T cells. Fig. 2D and 3C were performed in HeLa cells.

### Immunoprecipitations and Western blots

Immunoprecipitations of 3XF-PPIA and 3XF-Mutant PPIA transiently transfected cells were performed with a mouse monoclonal anti-FLAG antibody (clone M2, Millipore Sigma) in 293T cells. 3XF-Mutant PPIA (G104A mutant) has reduced catalytic activity due to blocked substrate access to the active site^23^. IP was performed on the cytoplasmic fraction of the cells. Western blots were done with a rat monoclonal anti-HA high-affinity antibody (clone 3F10, Millipore Sigma), a rabbit polyclonal anti-histone H3 antibody (ab1791, Abcam), and a rabbit polyclonal anti-cyclophilin A antibody (#2175, Cell Signaling Technology).

### Pulsed SILAC (Stable Isotope Labelling with Amino acids in Cell culture)

The workflow of the pulsed SILAC experiment performed in this study is described in Extended Data Fig. 8. First, control and PPIA knock-down (Kd) HeLa or 293T cells were cultured for five days in standard DMEM medium. Once cells have reached a similar confluence level (∼50%), heavy isotope (^13^C-^15^N-Lysine and ^13^C-^15^N-Arginine)-containing DMEM medium (Thermo Fisher Scientific) was added in excess to the cells for 24 h. Cells were harvested and 100 µg of protein cell lysates from each cell type and condition were subjected to acetone precipitation; subsequent denaturation, reduction, and alkylation prior to overnight in-solution digestion at 37°C with trypsin in order to generate peptides for mass spectrometry. Digestions were terminated by adding equal volume of 2% formic acid, and then desalted with Oasis HLB 1 ml reverse phase cartridges (Waters) according to the vendor’s procedure.

#### Liquid chromatography-tandem mass spectrometry (LC-MS/MS) analysis

An aliquot of the tryptic digest was analysed by LC-MS/MS on an Orbitrap Fusion Tribrid mass spectrometer (Thermo Scientific) interfaced with an UltiMate 3000 Binary RSLCnano System (Dionex), as previously described^54^. In our experiments, dynamic exclusion was employed for 40 s.

#### Data processing and analysis

The raw proteomic files were processed with the Proteome Discoverer 1.4 software (Thermo Scientific) and MS/MS spectra were searched against Uniprot-Homo sapiens database using the SEQUEST HT search engine. The spectra were also searched against decoy database using a peptide target false discovery rate (FDR) set to <1% and < 5%, for stringent and relaxed matches, respectively. The search parameters allowed for a maximum of two missed trypsin cleavages and set MS/MS tolerance to 0.6 Da. Carbamidomethylation on cysteine residues was used as fixed modification while oxidation of methionine as well as SILAC heavy arginine (^13^C_6_- ^15^N_4_), and SILAC heavy lysine (^13^C_6_-^15^N_2_) were set as variable modifications. Quantification of SILAC pairs was performed with the Proteome Discoverer software. Precursor ion elution profiles of heavy vs. light peptides were determined with a MS tolerance of 3 ppm. The area under the curve was used to determine a SILAC ratio for each peptide.

### Proximity Ligation Assay (PLA)

Whole bone marrow was obtained from the hind limb long bones and hip bones of young and old animals (5 month-old and 23 month-old, respectively). Lineage-positive cells were isolated using the Direct Lineage Cell depletion kit (Miltenyi Biotec) and magnetically depleted with an AutoMACS Pro Separator (Miltenyi Biotec). Then, the lineage-negative fraction was resuspended at a concentration of 10^8^ cells/ml and stained on ice for 15 min with the combination of antibodies characterizing HSCs described in the “Cell analysis and FACS” section above. Cell sorting was carried out on an Aria I FACS instrument (BD Biosciences). Finally, isolated HSCs were cytospinned, attached onto a Cellview slide (#543979, Greiner Bio-one), and fixed in 4% paraformaldehyde.

To quantify PPIA expression, proximity ligation assays were performed on isolated HSCs with the Duolink in Situ Red Starter Kit Mouse/Rabbit (DUO92101, Millipore Sigma), adapting the vendor’s protocol for HSCs. Briefly, HSCs were permeabilized with PBS + 0.5% Triton X- 100 for 7 min, washed with PBS, and blocked in 5% donkey serum for 30 min at room temperature. After a short wash in PBS, slides were incubated in a humidity chamber for 1 h at 37°C with Duolink blocking solution. Then, primary antibodies (mouse anti-cyclophilin A antibody, ab58114, and rabbit anti-cyclophilin A antibody, ab41684; both from Abcam) were applied overnight at 4°C in a humidity chamber. After washing the samples twice with Duolink buffer A, the diluted anti-mouse PLUS and anti-rabbit MINUS PLA probes were added to the samples for 1 h at 37°C in a pre-heated humidity chamber. Following two washes with buffer A, the cells were incubated with a DNA ligase previously diluted in Duolink Ligation buffer for 30 min at 37°C. Then, samples were washed twice in Duolink buffer A under gentle shaking and incubated with a diluted DNA polymerase solution for 1 h 40 min at 37°C in the dark. Finally, slides were rinsed twice in 1X wash buffer B for 10 min and once in 0.01X wash buffer B for 1 min at room temperature and mounted with Duolink *in situ* mounting medium containing DAPI. For each PPIA antibody, a negative control experiment was performed where only one antibody or no antibody was incubated with the PLA probes (data not shown). Fluorescence was visualized with a Celldiscoverer7 confocal microscope (Zeiss) at 100X magnification and images were processed for background subtraction and orthogonal projection with the ZEN Pro imaging software (Zeiss). The experimenter was blinded to the origin of the samples during the PLA staining and spot counting. An average of 90 cells per condition was counted.

### RNA-sequencing (RNA-seq)

#### Isolation and selection of HSPCs

Wild-type HSPCs were isolated from the hind limb long bones of 4 to 6 month-old and 31 to 33 month-old male mice. c-Kit^+^ cells were stained and magnetically isolated from the lineage-depleted cell suspension using the EasySep mouse CD117 (c-Kit) positive selection kit (Stem Cell Technologies), following the manufacturer’s instructions. After overnight growth in serum-free medium (StemSpan SFEM, Stem Cell Technologies), supplemented with murine TPO (20 ng/ml, PeproTech), SCF (10 ng/ml, PeproTech), and the beta-Catenin agonist CHIR99021 (250 nM, Stemgent), HSPCs were harvested as cell pellets. Immediately after, RNA extraction was carried out with the RNeasy Plus Mini kit with genomic DNA Eliminator columns (Qiagen) in combination with on-column DNaseI digestion (Qiagen), according to the vendor’s protocol.

#### Preparation and sequencing of RNA-seq libraries

Total RNA-seq libraries were generated and prepared for multiplexing on the Illumina platform with the TruSeq stranded total RNA library prep (Illumina) according to manufacturer’s protocol. Libraries included ERCC ExFold RNA spike-in mixes (Thermo Fisher Scientific) to assess the platform dynamic range. RNA spike-in mixes confirmed high fidelity between two independent NGS runs (R^2^= 0.991 and 0.943, respectively) (SI-Mouse HSPCs RNA seq). The resultant libraries were quality-checked on a Bioanalyzer 2100 instrument (Agilent) and quantified with a Qubit fluorometer (Thermo Fisher Scientific). Further quantification of the adapter ligated fragments and confirmation of successful P5 and P7 adapter incorporations were assessed with the KAPA universal library quantification kit for Illumina (Roche) run on a ViiA7 real-time PCR system (Applied Biosystems). Multiplexed and equimolarly pooled library products were re-evaluated on the Bioanalyzer 2100 and diluted to 18 pM for cluster generation by bridge amplification on the cBot system. Then, libraries were loaded onto a HiSeq2500 rapid run mode flowcell v2, followed by paired-end 100 cycle sequencing run on a HiSeq2500 instrument (Illumina). PhiX Control v3 adapter-ligated library (Illumina) was spiked-in at 2% by weight to ensure balanced diversity and to monitor clustering and sequencing performance. We obtained a minimum of 50 million reads per sample.

#### Data processing

Fastq file generation was achieved with the Illumina’s BaseSpace Sequence Hub. Demultiplexing was based on sample-specific barcodes. All bioinformatic analyses were performed with Linux command line tools. After removing the short sequence reads which did not pass quality control and discarding reads containing adaptor sequences with Cutadapt v.1.12^55^, sequence reads were assembled and mapped against the mouse MM9 reference genome (Genome Reference Consortium) with TopHat2/Bowtie2 v.2.1.0^56^. Gene expression changes were quantified with Cufflinks and Cuffdiff v.2.1.1^57^ and data were normalized by calculating the fragments per kilobase per million mapped reads (FPKM).

### Statistics

All statistical analyses were performed using Stata v.12, Stata v.15.1, or GraphPad Prism 8 software. Statistical analysis for individual gene analyses and transplantation data was performed using a two-tailed Student’s *t*-test while large datasets were compared with a two-sided Wilcoxon rank-sum test or a Chi square test with Yates’ correction. On violin plots, the dashed line marks the median and the dotted lines represent the lower and upper quartiles.

## Data Availability

Mass spectrometry data obtained after 3XF-PPIA immunoprecipitation (Fig. 2A) are deposited with the ProteomeXchange Consortium (http://proteomecentral.proteomexchange.org/) via the MASSIVE repository (MSV000083867) with the dataset identifier PXD014025.

The datasets generated in the mouse HSC proteome profiling (Fig. 4B) have been deposited to the ProteomeXchange Consortium via the MASSIVE repository (MSV000083845) with the dataset identifier PDX013995. For human HSPC/lymphocyte/granulocyte proteome and transcriptome, results from a previously published report were analysed^47^. HSPC data analysis was based on levels of mRNA and proteins belonging to age-affected pathways in individual HSPC population. “Pathways were required to have between 5 and 150 members to be sufficiently covered and to have at least 20% of its quantified components being significantly altered upon aging”, as defined by the original authors. Lymphocyte and granulocyte peptide comparison was based on the percentage of IDRs per protein for polypeptides that are unique to lymphocytes or unique to granulocytes. Rare peptides that were identified in less than 50% of the pooled lymphocyte and the pooled granulocyte analyses were excluded.

For transcriptomic analysis of murine HSPCs (SI-Mouse HSPCs RNA seq), raw and processed RNA-seq data have been deposited with the Gene Expression Omnibus (GEO) database under accession code GSE151125.

## Acknowledgements

A.C. was supported by the Cancer Prevention and Research Institute of Texas (CPRIT- RR140038), the Ted Nash Long Life Foundation, and NIH R01DK115454. We would like to thank Dr. Margaret Goodell and Dr. Joanne Ino Hsu for helpful discussions, Catherine Gillespie for editorial assistance, and Laura Prickett-Rice and Kat Folz-Donahue for FACS support.

We acknowledge the Genomic and RNA Profiling Core (supported by NIH-NIDDK P30DK56338 Center grant, NIH-NCI P30CA125123 Center grant, and NIH 1S10OD02346901 S10 grant), the Cytometry and Cell Sorting Core (supported by CPRIT Core Facility Support Award (CPRIT-RP180672) and NIH (CA125123 & RR024574) grants), and the Mass Spectrometry Proteomics Core (supported by NIH-NCI P30CA125123 Center grant and CPRIT Core Facility Support Award (CPRIT-RP170005)) at Baylor College of Medicine for their technical support. We thank Li Li and Dr. Sheng Pan at the Clinical and Translational Proteomics Service Center at the University of Texas Health Science Center for assistance with the generation and analysis of the pulsed SILAC MS data.

## Author Contributions and Competing Interest Declaration

L.M. designed and conducted molecular assays, interpreted results, and wrote the manuscript. P.I. conducted molecular assays, interpreted results, and contributed to manuscript preparation. S.E.M. helped with the acquisition of microscopy images. D.B.S., C.T.H., D.S.K., and B.S. helped with *in vitro* assays and functional haematopoietic assays and edited the manuscript. E.S. (Whitehead Institute) and J.C.K. were involved in protein identification. B.C.B. supported the *in vivo* studies, discussions, interpreted data, and edited the manuscript. E.S. (Baylor College of Medicine) supported data analysis and edited the manuscript. D.T.S. co- supervised the haematopoietic aspects of this study, interpreted data, and edited the manuscript. A.C. supervised this study and was involved in all experimental aspects, conceived the project, analysed data, and wrote the manuscript.

Supplementary Information is available for this paper. Correspondence and requests for materials should be addressed to A.C. The authors declare no competing financial interests.

## Extended Data Figure Legends

**Extended Data Fig.1:**
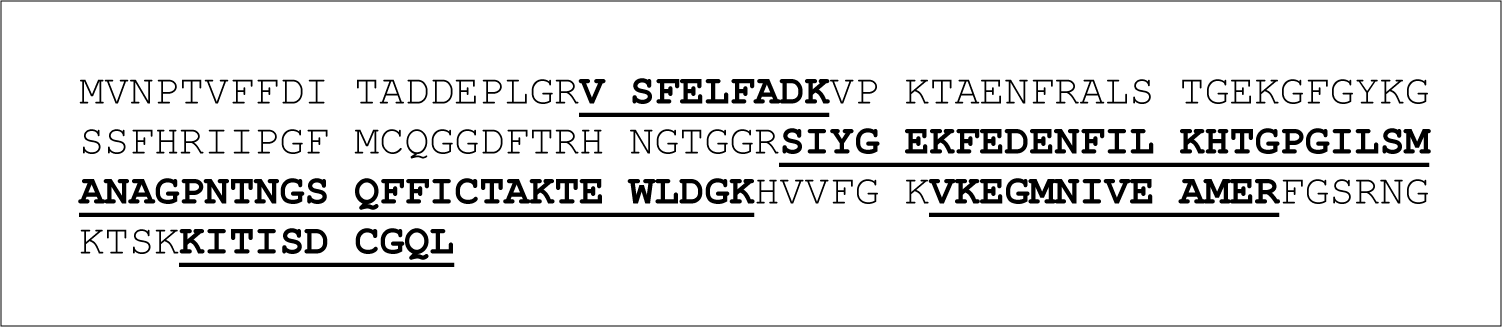
Identification coverage of PPIA. Spots were excised from 2-D SDS PAGE for extraction and trypsin digestion. MS/MS spectra cover 49.8% of the entire PPIA protein sequence (underlined).

**Extended Data Fig.2:**
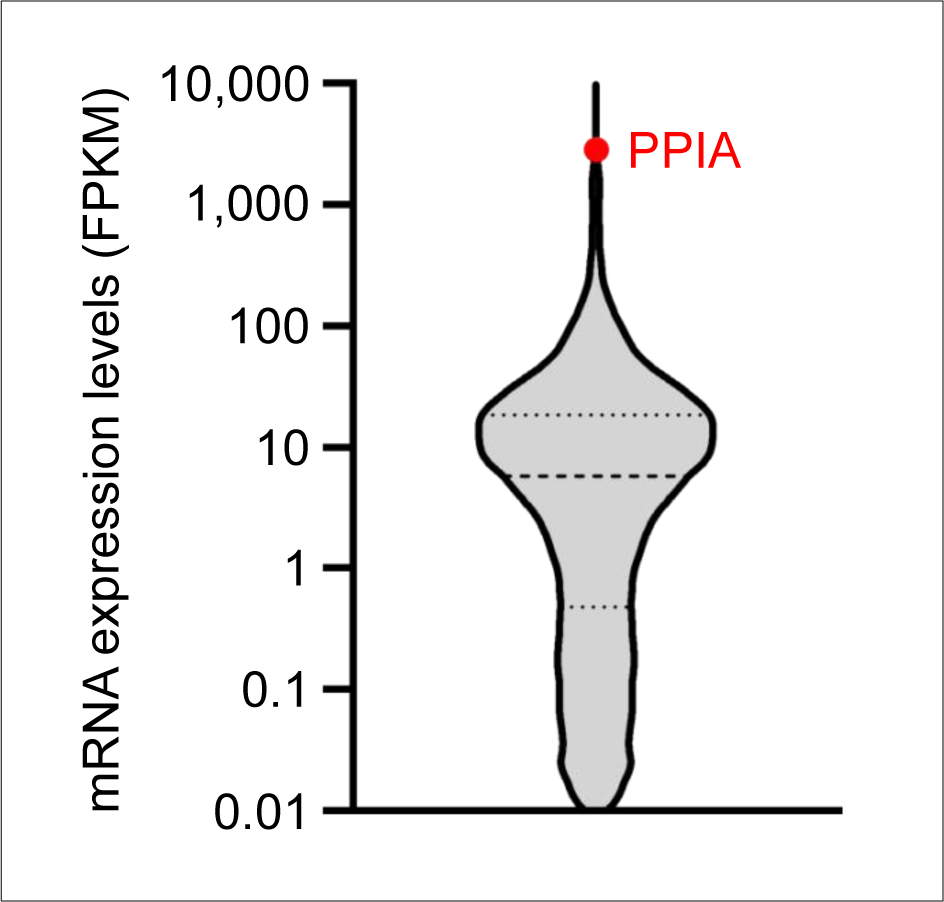
PPIA is the most highly transcribed chaperone gene in HSPCs. Violin plot represents the per gene distribution of RNAseq reads in the mouse HSPC transcriptome. PPIA is the 6^th^ most highly expressed gene out of 15,778 genes. Red spot indicates PPIA transcript.

**Extended Data Fig.3:**
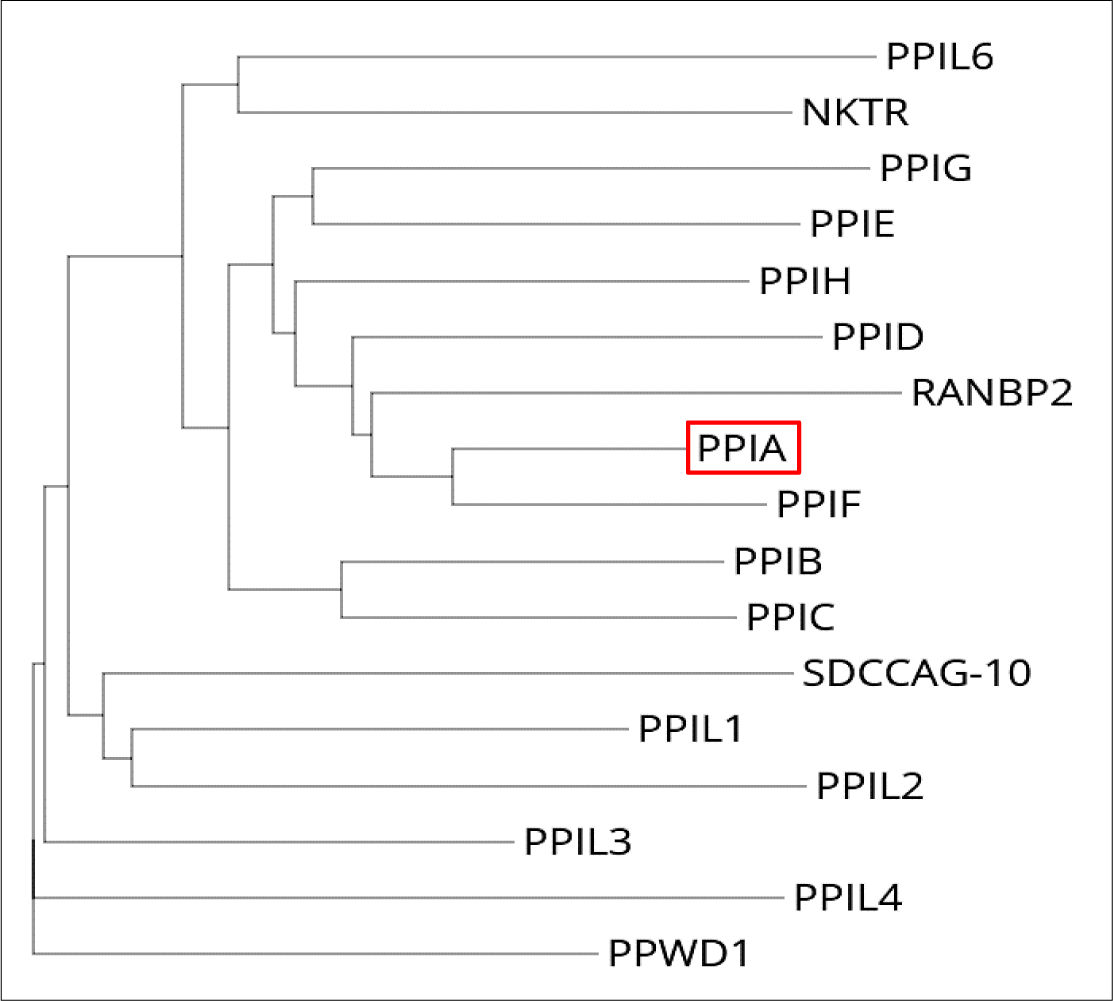
Phylogenetic tree of the Cyclophilin protein family in humans. The tree was based on protein sequence alignments from the NCBI RefSeq database with the EMBL Clustal Omega program. The red box indicates PPIA.

**Extended Data Fig.4:**
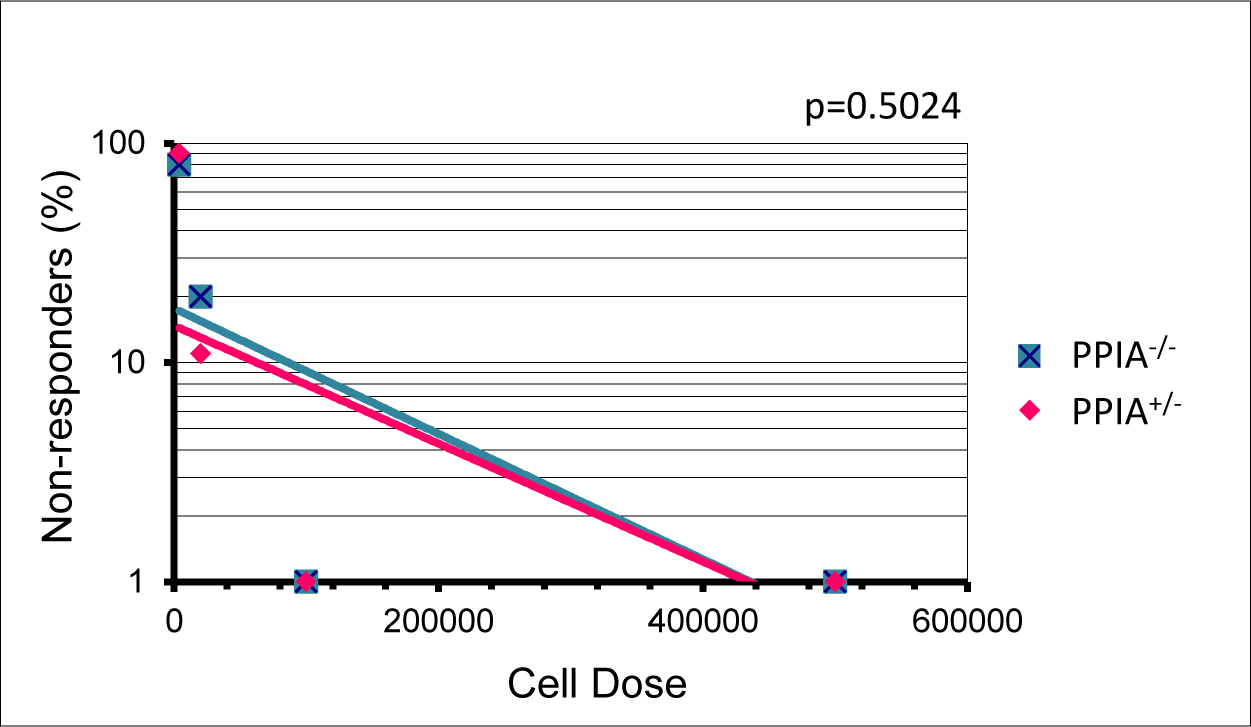
Limiting Dilution Transplantations of PPIA heterozygous (PPIA^+/-^) and knockout (PPIA^-/-^) bone marrow. 500,000 competitor cells (CD45.1^+^) were co-injected with 4,000, 20,000, 100,000, or 500,000 nucleated bone marrow cells of PPIA^+/-^ or PPIA^-/-^ mice. Reconstitution of peripheral CD45.2^+^ cells was assayed 20 weeks after transplantation and differences were compared using a two-tailed Poisson t-test. No significant difference exists between PPIA heterozygous and deficient donors.

**Extended Data Fig. 5:**
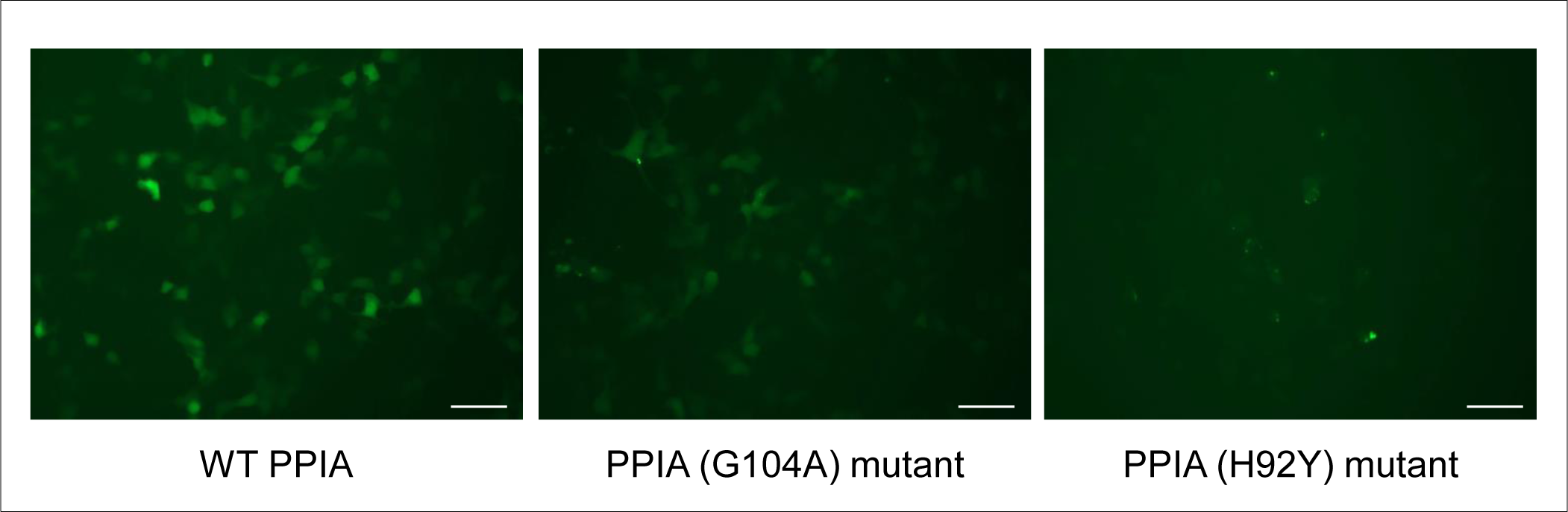
Expression pattern of wild-type (WT) and mutant PPIA proteins. 293T cells were transiently transfected with WT PPIA-GFP, PPIA(G104A)-GFP mutant or PPIA(H92Y)-GFP catalytic core mutant. The expression pattern of the WT and PPIA mutant proteins was assessed with an EVOS cell imaging system, using a GFP filter. Scale bar=50 µm.

**Extended Data Fig. 6:**
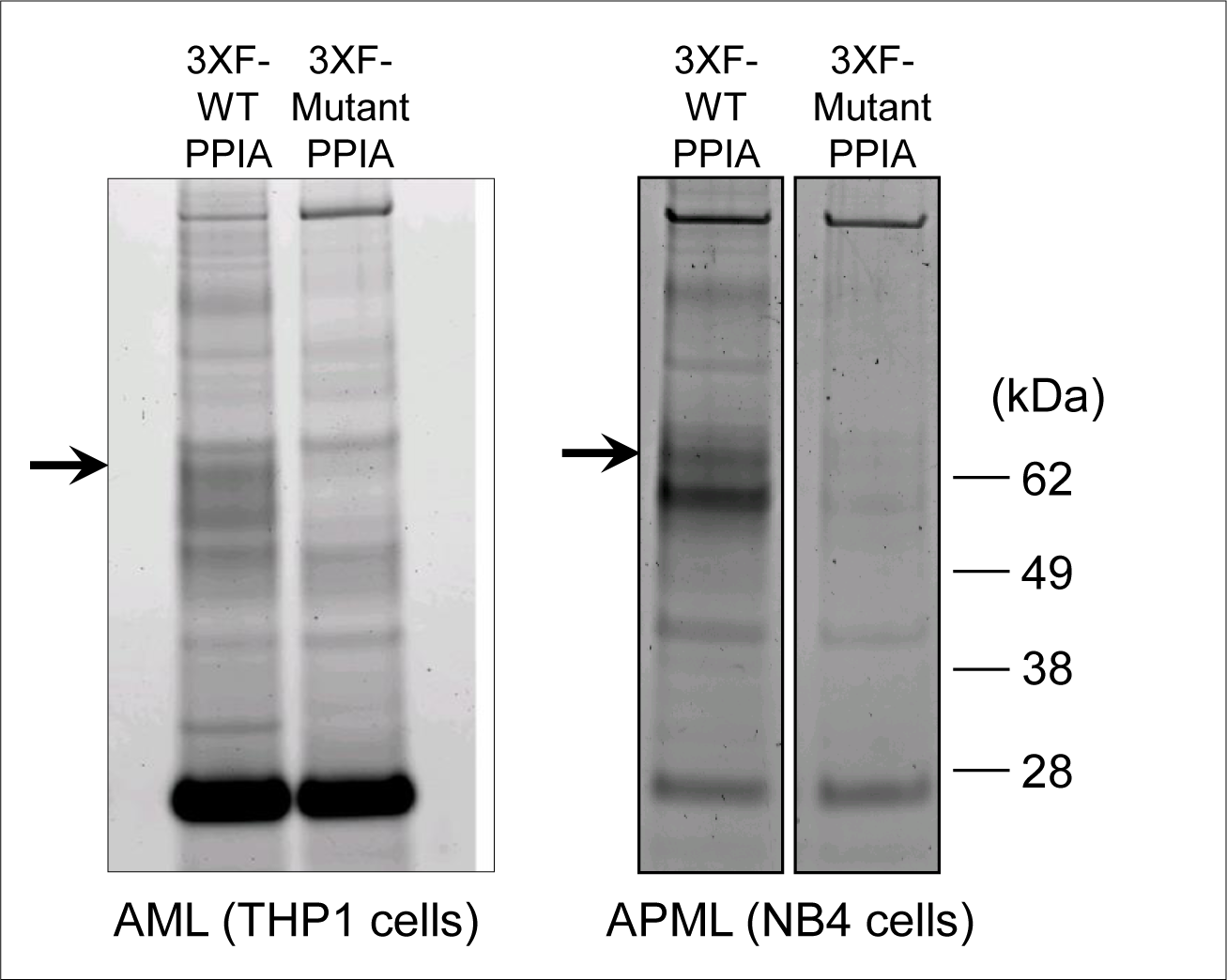
Interaction between PPIA and PABPC1 in haematopoietic cells. Co-IP followed by MS/MS identifies PABPC1 as an interactor of PPIA in the human haematopoietic cell lines THP1 (acute monocytic leukemia, AML) and NB4 (acute promyelocytic leukemia, APML). Cells were transduced with 3XF-tagged PPIA (3XF-WT-PPIA or 3XF-Mutant(G104A)-PPIA, respectively) and IP was performed with an anti-3XFLAG antibody. The arrow indicates the band for PABPC1 protein.

**Extended Data Fig. 7:**
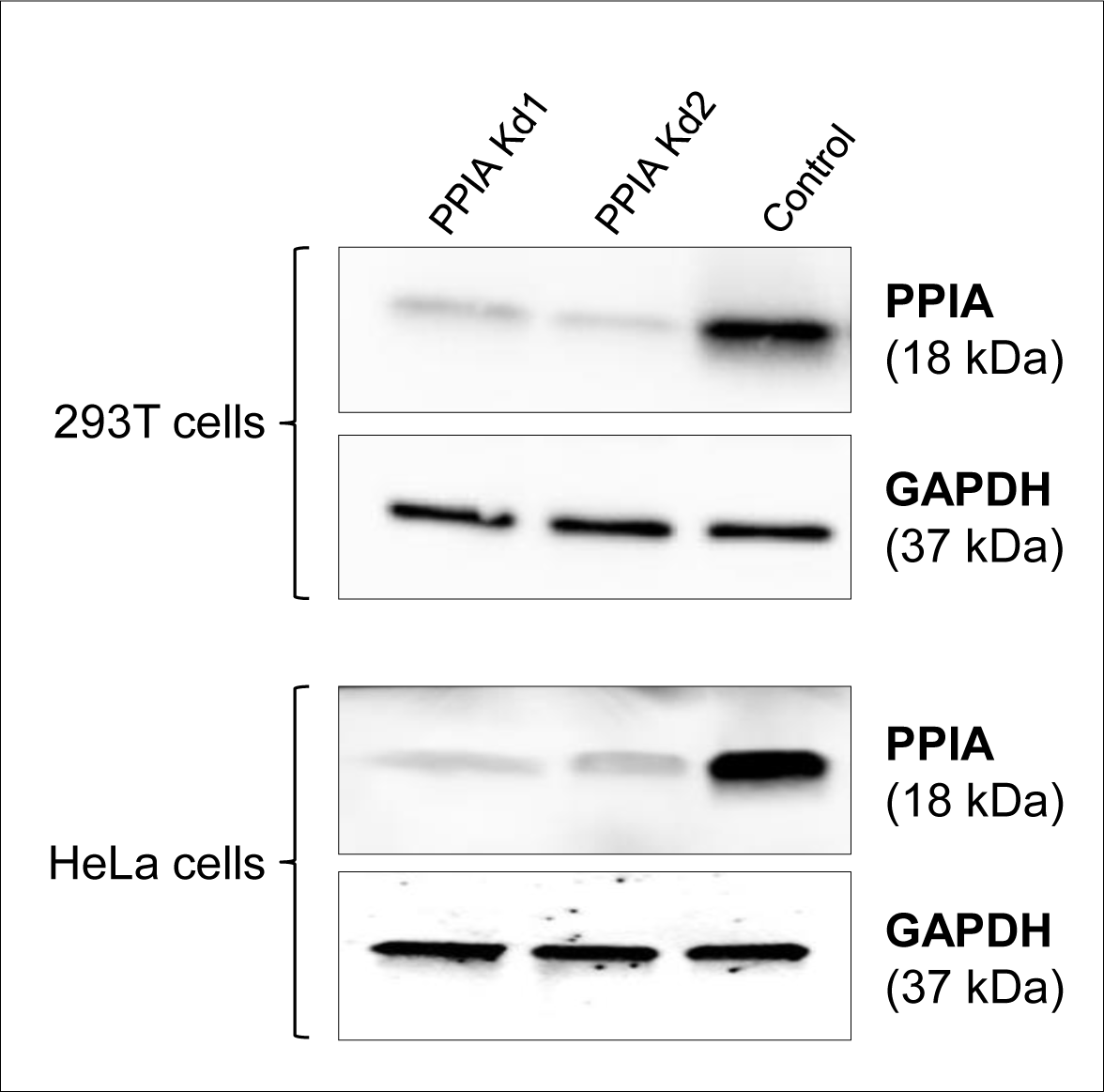
Efficiency of PPIA knockdown in 293T cells and HeLa cells. Cells were stably transduced with pLKO.1-TRC control, TRC PPIA Kd1, or TRC PPIA Kd2 lentiviral vectors, respectively. Then, cell lysates were prepared and loaded onto a SDS-PAGE in order to measure PPIA protein expression by Western-blot using a rabbit polyclonal anti-PPIA antibody. Glyceraldehyde 3-phosphate dehydrogenase (GAPDH) was used as a loading control for protein normalization.

**Extended Data Fig. 8:**
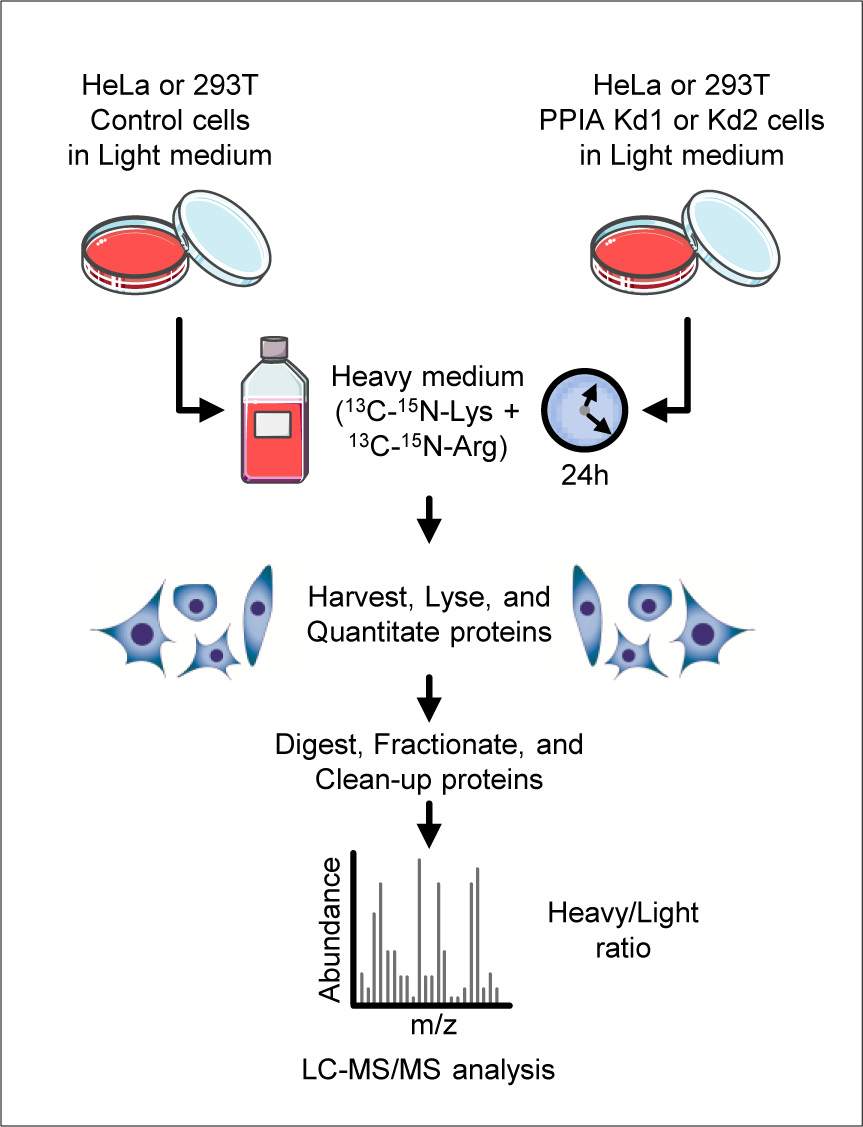
Schematic representation of the pulsed SILAC (stable-isotope labelling in cell culture) experimental design used to evaluate protein *de novo* synthesis in this study. Following five days of growth in standard DMEM medium (containing light/unlabeled variants of lysine and arginine), heavy lysine and arginine (^13^C-^15^N-Lys and ^13^C-^15^N-Arg) were added in excess to control or PPIA knock-down (Kd) HeLa or 293T cells. After medium exchange, newly synthesized proteins incorporate the heavy label while pre-existing proteins remain unlabeled. 24 h later, cells were harvested and protein lysates were digested. Pulse treatment for 24h allowed for metabolic labeling of newly translated proteins. The resulting peptides from each cell type and condition were submitted separately to liquid chromatography-tandem mass spectrometry (LC-MS/MS) analysis. Protein *de novo* synthesis was determined by the heavy to unlabeled ratio quantified by mass spectrometry, as previously described^27, 58^.

**Extended Data Fig. 9:**
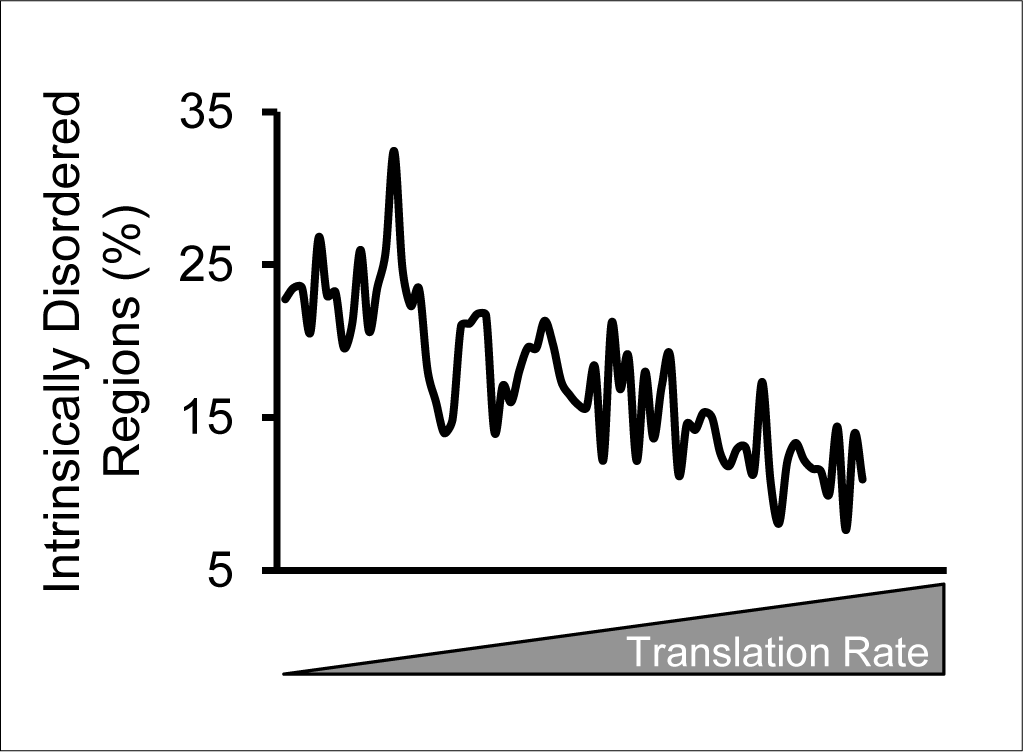
Presence of intrinsically disordered regions correlates with a slower translation rate. Data analysis showing the inverse correlation between protein translation speed^27^ and percentage of intrinsically disordered regions^28^ in the whole proteome. Proteins with more intrinsically disordered regions translate at a slower rate than structured proteins.

**Extended Data Fig. 10:**
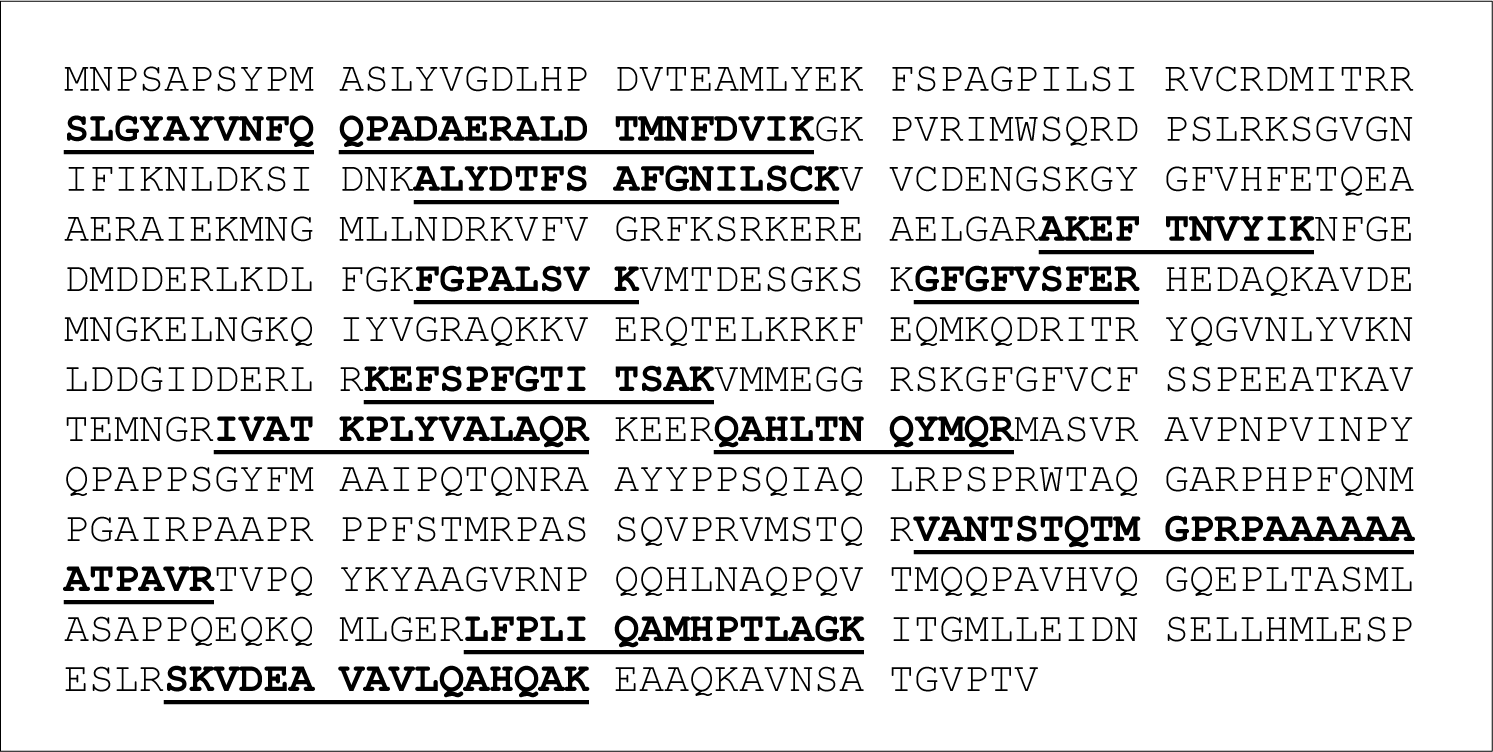
Identification coverage of PABPC1. Co-immunoprecipitated bands were excised after SDS-PAGE (Fig. 2A), trypsin digested, and identified by MS/MS. Coverage for PABPC1 extends to 26.4% of the protein, including the N-terminal RNA-binding domain, the C-terminal cap-binding domain, and the unstructured linker region.

**Extended Data Fig. 11:**
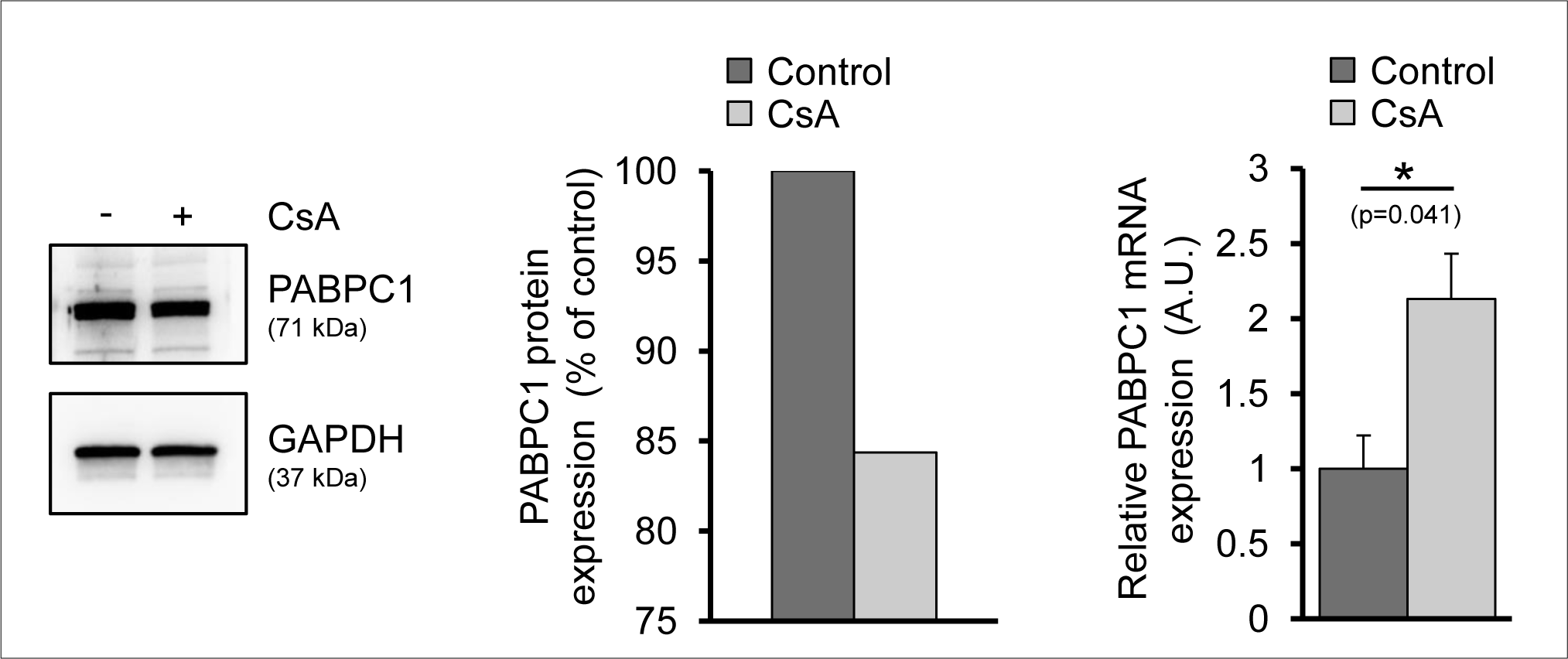
PABPC1 protein, but not transcript levels, are slightly decreased upon PPIA inhibition by cyclosporine A in MM.1S cells. Cyclosporine A (CsA, 10 µM, Millipore Sigma) treatment was performed for 48 hours with DMSO as solvent control. Multiple myeloma cells were chosen for their high level of protein synthesis due to production of monoclonal immunoglobulins. Western blot image is representative of two independent biological replicates. RT-qPCR data are mean ± SD of two independent experiments performed with triplicate samples; *p < 0.05 by unpaired Student’s two-tailed t-test. A.U., Arbitrary Unit.

**Extended Data Table 1:**
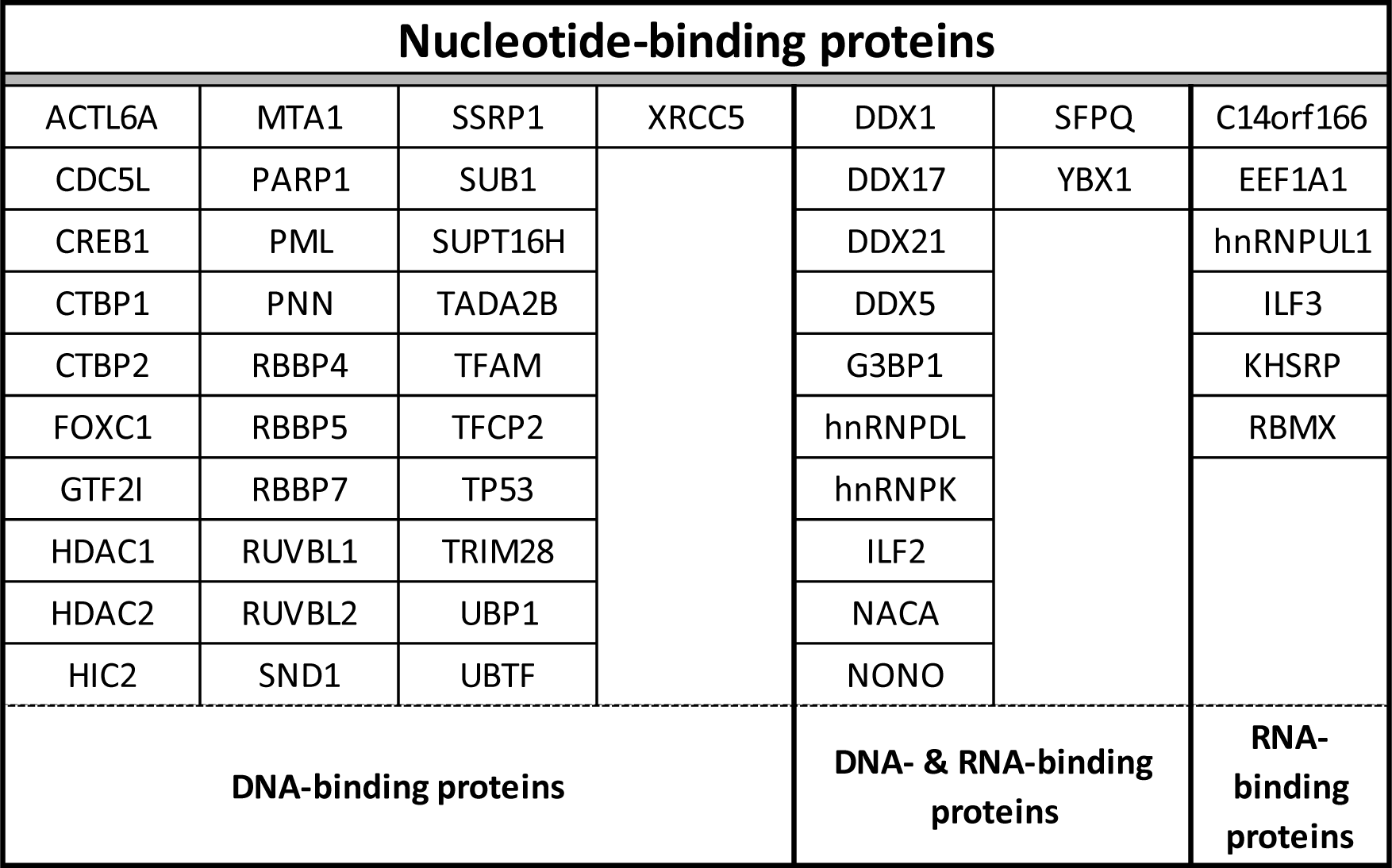
List of PPIA client proteins which are nucleotide-binding proteins. Proteins are listed by their official gene name.

**Extended Data Table 2:**
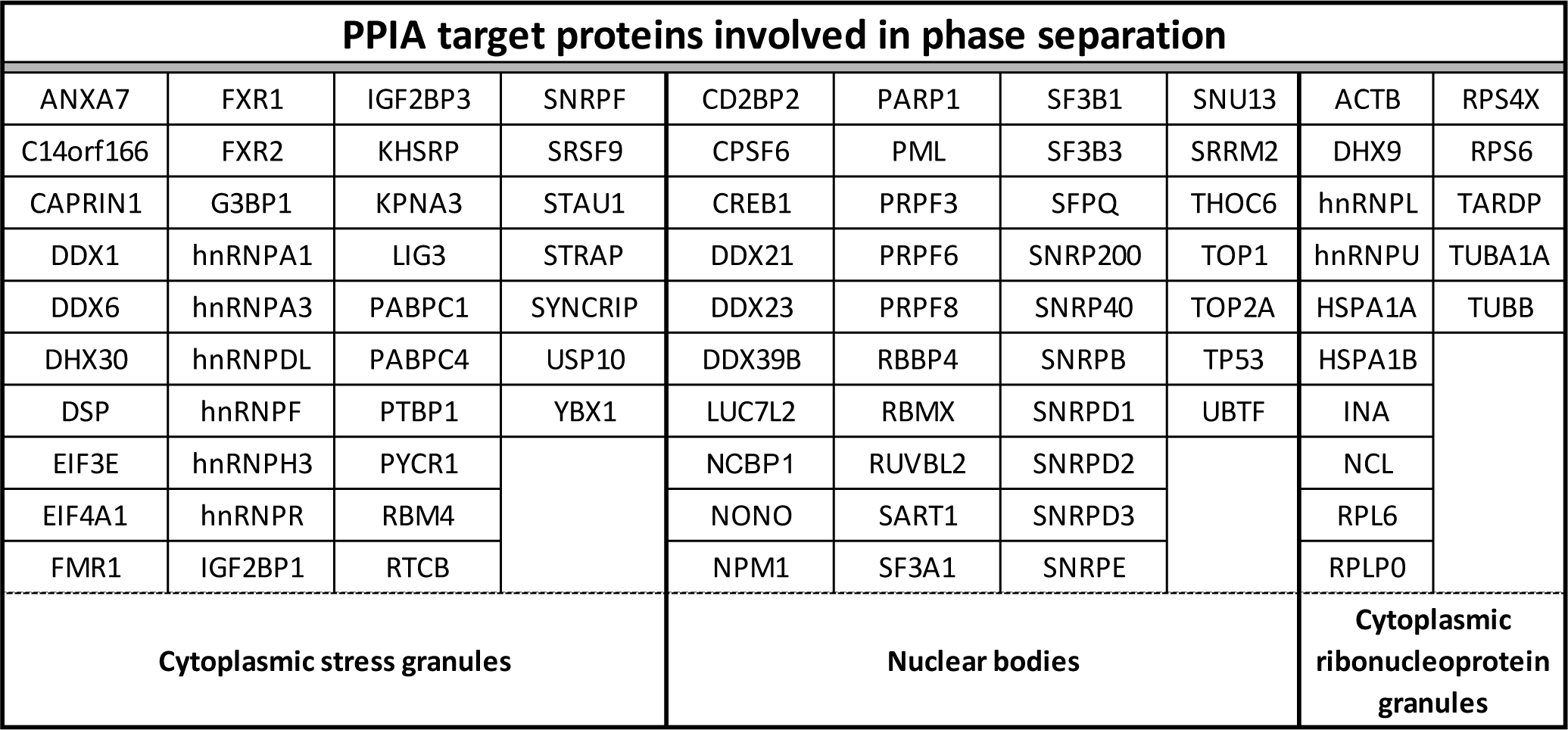
List of PPIA client proteins which are involved in protein phase separation. Proteins are listed by their official gene name. Nuclear bodies include nucleoli, Cajal bodies, nuclear speckles, paraspeckles, PML nuclear bodies, and histone locus bodies.

## Notes

### Competing Interest Statement

The authors have declared no competing interest.

